# Genomic rearrangements generate hypervariable mini-chromosomes in host-specific lineages of the blast fungus

**DOI:** 10.1101/2020.01.10.901983

**Authors:** Thorsten Langner, Adeline Harant, Luis B. Gomez-Luciano, Ram K. Shrestha, Joe Win, Sophien Kamoun

## Abstract

Supernumerary mini-chromosomes–a unique type of genomic structural variation–have been implicated in the emergence of virulence traits in plant pathogenic fungi. However, the mechanisms that facilitate the emergence and maintenance of mini-chromosomes across fungi remain poorly understood. In the blast fungus *Magnaporthe oryzae*, mini-chromosomes have been first described in the early 1990s but, until very recently, have been overlooked in genomic studies. Here we investigated structural variation in four isolates of the blast fungus *M. oryzae* from different grass hosts and analyzed the sequences of mini-chromosomes in the rice, foxtail millet and goosegrass isolates. The mini-chromosomes of these isolates turned out to be highly diverse with distinct sequence composition. They are enriched in repetitive elements and have lower gene density than core-chromosomes. We identified several virulence-related genes in the mini-chromosome of the rice isolate, including the polyketide synthase *Ace1* and the effector gene *AVR-Pik*. Macrosynteny analyses around these loci revealed structural rearrangements, including inter-chromosomal translocations between core- and mini-chromosomes. Our findings provide evidence that mini-chromosomes independently emerge from structural rearrangements of core-chromosomes and might contribute to adaptive evolution of the blast fungus.

**Author summary:** The genomes of plant pathogens often exhibit an architecture that facilitates high rates of dynamic rearrangements and genetic diversification in virulence associated regions. These regions, which tend to be gene sparse and repeat rich, are thought to serve as a cradle for adaptive evolution. Supernumerary chromosomes, i.e. chromosomes that are only present in some but not all individuals of a species, are a special type of structural variation that have been observed in plants, animals, and fungi. Here we identified and studied supernumerary mini-chromosomes in the blast fungus *Magnaporthe oryzae*, a pathogen that causes some of the most destructive plant diseases. We found that rice, foxtail millet and goosegrass isolates of this pathogen contain mini-chromosomes with distinct sequence composition. All mini-chromosomes are rich in repetitive genetic elements and have lower gene densities than core-chromosomes. Further, we identified virulence-related genes on the mini-chromosome of the rice isolate. We observed large-scale genomic rearrangements around these loci, indicative of a role of mini-chromosomes in facilitating genome dynamics. Taken together, our results indicate that mini-chromosomes facilitate genome rearrangements and possibly adaptive evolution of the blast fungus.

## Introduction

Genomes of plant pathogens are highly dynamic and typically exhibit an architecture that facilitates rapid adaptation to their hosts. Since the rise of genome sequencing it became evident that plant pathogen genomes are often structured to accommodate high genetic diversification rates at virulence-related loci while maintaining relative stability in house-keeping regions, a phenomenon that shaped the term “two-speed genome” [1]. Since then, the two-speed genome concept has been widely documented in a number of plant pathogenic species [2,3]. Interestingly, various types of genome architecture have been observed in different species. These include effector gene clusters [4,5], lineage-specific genomic regions that are rich in transposable elements [6–10], or enrichment of virulence related genes in specific genomic regions, e.g. unstable telomeric and sub-telomeric regions [11]. Typically, these genomic compartments display higher rates of adaptive mutations compared to the rest of the genome [12]. In addition to signatures of single nucleotide polymorphisms (SNPs) indicative of positive selection and presence/absence polymorphisms, structural variation is common in pathogen populations and ranges from copy number variations of single genes to chromosome-scale rearrangements [13,14]. Extreme cases of genomic rearrangement are large-scale, chromosome length variations and the presence of isolate-specific, supernumerary chromosomes (syn. B-, accessory-, conditionally dispensable, mini-chromosomes). These are usually small, non-essential chromosomes that occur in addition to the regular set of conserved chromosomes within a species and have been described in animals, plants, and fungi [14,15]. Supernumerary chromosomes are present at different frequencies in natural populations of eukaryotic plant pathogens [14,16]. In some fungal species, supernumerary chromosomes have been directly implicated in the emergence of new virulence traits, underpinning the importance of understanding their role in evolution and pathogen adaptation [17–19]. However, the diversity of supernumerary chromosomes across plant pathogens and their contribution to genome plasticity is still poorly known.

Supernumerary chromosomes share common features that distinguish them from core-chromosomes. Although they are variable in size, they tend to be smaller than core-chromosomes with their size ranging from ∼400 kb to 3 Mb in plant pathogenic fungi [8,14]. Yet, despite their small size, supernumerary chromosomes can contribute up to 15% to the total genome in certain species. Their number is also variable with some plant pathogenic fungi containing up to eight supernumerary chromosomes in addition to the core-chromosome set [9]. Dynamic loss or gain of supernumerary chromosomes has been observed especially under stress conditions indicating that supernumerary chromosomes are major drivers of genome plasticity [20]. In addition to frequent genomic rearrangements, supernumerary chromosomes often do not follow Mendelian inheritance. They tend to be meiotically unstable, and thus, frequently lost [21]. However, in some cases meiotic gene drives have been observed increasing their potential to be inherited and potentially explaining their abundance in pathogen populations [22].

Supernumerary chromosomes are usually gene poor and repeat rich compared to the core-chromosomes, and form one illustration of the two-speed genome concept [12,23]. However, a clear association with adaptive evolution is not always evident as they do not necessarily carry virulence-related genes [9,24]. Nonetheless, in some plant pathogenic fungi, supernumerary chromosomes are directly implicated in virulence [17,18]. The origin of supernumerary chromosomes is still debated but there is evidence for segmental duplications from core-chromosomes (or ancient core-chromosome duplication followed by partial chromosome loss) and horizontal chromosome transfer [19,25–27]. Interestingly, supernumerary chromosomes can be transferred between isolates independently of the core genome and can alter the virulence spectrum of plant pathogens [19,27]. It is thus possible that supernumerary chromosomes facilitate gene flow in natural populations leading to new pathotypes.

Plant pathogens are ubiquitous in the environment and can cause severe damage to both cultivated and wild plant species [28–31]. Plant pathogens are often specialized on a specific host species or taxon. At the center of the co-evolutionary dynamics between pathogens and plants are effector proteins, i.e. secreted proteins that manipulate host processes to facilitate infection and colonization [32–35].In return, host plants have evolved immune receptors that can detect conserved molecular patterns and effector proteins to defend against invading pathogens [36,37]. This generally leads to fast-paced co-evolution between pathogens and their host plants that often follows arms-race dynamics where the frequency of adaptive mutations rises quickly in pathogen populations [16].

The blast fungus *M. oryzae* (Syn. *Pyricularia oryzae*) causes blast disease, one of the most devastating crop diseases worldwide resulting in yield losses in rice and wheat that make it a threat to global food security [28,29,38,39]. Despite its Linnaean binomial name, *M. oryzae* is a multihost pathogen that can infect more than 50 cultivated and wild grass species. Population genomics of *M. oryzae* revealed that the species is formed by an assemblage of differentiated lineages that are associated with particular host taxa, such as important cereals like rice (*Oryza sativa*), finger millet (*Eleusine coracana*), wheat (*Triticum aestivum*), and foxtail millet (*Setaria italica*), as well as weeds such as Indian goosegrass (*Eleusine indica*) [40,41]. Records of rice blast disease in China date back to the early 17^th^ century and until today it is recognized as one of the most threatening and widely distributed rice diseases [39]. Recent population genetics studies revealed that the rice-infecting lineage of *M. oryzae* consists of both a recombining population and multiple, clonally expanded lineages [42–44]. In Europe, the rice blast fungus population consists of only one of three major clonal lineages and it is possible that mating type isolation led to local asexual expansions. Another impactful blast disease is wheat blast. In the mid 1980s the disease emerged on wheat plants in Brazil and has since spread across large regions of South America and, more recently, South Asia, demonstrating the ability of *M. oryzae* to rapidly undergo host-range expansions and global pandemics [45–47].

Although signatures of gene flow have been observed within and between lineages [41], *M. oryzae* is thought to predominantly propagate asexually in agricultural settings. Given that genetically differentiated lineages tend to be associated with particular host genera, selection pressure imposed by the host plant is probably the main driver of adaptive evolution [41]. Adaptation to a specific host can be accompanied by gain or loss of effector genes that define the pathogen host range highlighting the importance of structural genomic variation [35,48–50]. The degree to which genome architecture facilitates structural variation and even gene flow is a fascinating and still poorly understood question [6,11,23,51].

*M. oryzae* mini-chromosomes have first been described in the early 1990s, when studies on karyotype diversity revealed that large chromosomal rearrangements occur frequently within and between clonal lineages [52,53]. More recently, Chuma *et al.* [11] demonstrated that the effector gene *AVR-Pita* underwent multiple translocation events and proposed that genomic location of effector genes to subtelomeric regions could favor rearrangements within the genome and gene flow within asexual populations. Interestingly, *AVR-Pita* genes occur on different chromosomes, including supernumerary mini-chromosomes. Additionally, Luo *et al.* [54] and Kusaba *et al.* [55] showed that two variants of the AVR-Pik effector gene are present on a 1.6 Mb mini-chromosome in the Japanese rice blast isolate 84R-62B and that parts of this mini-chromosome can translocate to core-chromosomes in crosses. Further, loss of the mini-chromosome resulted in gain of virulence on host plants carrying the rice immune receptor Pik, a phenotype due to the associated loss of the *AVR-Pik* genes [55].

Despite the fact that *M. oryzae* mini-chromosomes have been known for 30 years, genome sequencing projects have somehow overlooked them. This has changed recently. Peng *et al.* [56] reported the first sequences of mini-chromosomes of *M. oryzae*. They analyzed the karyotypes of the three wheat blast isolates T25, B71, and P3. T25 was collected in Brazil in 1988 and B71 and P3 were collected in 2012 in Brazil and Paraguay, respectively. B71 contains a 2 Mb mini-chromosome and P3 contains two mini-chromosomes of 1.5 and 3 Mb, whereas T25 does not contain any mini-chromosomes. The mini-chromosomes of B71 and P3 only share partial sequence similarity, have lower gene and higher repeat content, and display partial similarity to sub-telomeric regions of core-chromosomes. The mini-chromosome of isolate B71 contains the effector genes *Pwl2* and *Bas1* in close proximity while they are located on separate core-chromosomes in other *M. oryzae* isolates. This raised the hypothesis that mini-chromosomes are sites of structural rearrangements associated with virulence factors within blast genomes. However, the genetic diversity of mini-chromosomes in other lineages of *M. oryzae*, and their association with genomic rearrangements and effector diversification remains poorly understood.

The objective of this study was to gain an overall picture of genomic structural variation across lineages of *M. oryzae* using the host-specific isolates from rice, foxtail millet, goosegrass and wheat that were previously sequenced using Illumina short reads by Chiapello et al. [48]. Our analyses led us to focus on mini-chromosomes, which we detected in three of the examined isolates of *M. oryzae*. We used long and short read sequencing data in combination with mini-chromosome isolation sequencing (MCIS) to improve the previous genome assemblies [48] to near chromosome quality. We found that the sequence composition of mini-chromosomes is highly variable indicating independent emergence of individual mini-chromosomes during *M. oryzae* lineage evolution. Further, we identified effector and virulence-related genes in the mini-chromosome of the rice-infecting isolate FR13 although this protein class was not generally enriched in the three sequenced mini-chromosomes compared to the core chromosomes. We documented several structural rearrangements around virulence-related loci which raises the possibility that chromosome-scale variation, notably mini-chromosome genesis, plays a role in driving adaptive genome plasticity in the blast fungus.

## Results

### Near chromosome quality genome assemblies of four host-specific isolates of *M. oryzae*

Following emerging evidence underpinning the abundance of structural, genomic variation in plant pathogens we re-examined 4 previously sequenced *M. oryzae* genomes of the host specific strains FR13 (rice), US71 (foxtail millet), CD156 (goosegrass), and BR32 (wheat). We used long read sequencing to generate highly contiguous assemblies [57] and improved accuracy of the assemblies by applying a polishing pipeline using nanopore and published Illumina raw reads of the same strains [48]. To assess the completeness and quality of our assemblies we performed a benchmarking universal single-copy orthologs (BUSCO) analysis using the Sordariomycota database (https://busco.ezlab.org/) which confirmed 98 – 98.2 % completeness, similar to the chromosome quality reference genome of strain 70-15 (98.2 %; Table 1). Overall, the new assemblies have vastly improved contiguity with contig numbers reduced from 111-2,051 in previous assemblies to 17-55 in the new assemblies (Table 1). Moreover, we increased the proportion of well-assembled repeat rich regions, reduced the number of ambiguous bases (“Ns”) to zero, and improved overall completeness (Table 1). These highly contiguous assemblies can thus serve as new reference genomes for host-specific isolates.

**Table 1:** Comparative summary statistics of genome assemblies.

| Host Isolate | *Oryza sativa* |  | *Setaria italica* |  | *Eleusine indica* |  | *Triticum sp.* |  | *Oryza sativa* |
| --- | --- | --- | --- | --- | --- | --- | --- | --- | --- |
|  | FR13 GEMO* | FR13 | US71 GEMO* | US71 | CD156 GEMO* | CD156 | BR32 GEMO* | BR32 | 70-15 |
| Technology | Illumina / 454 | ON-MinION | Illumina / 454 | ON-MinION | Illumina / 454 | ON-MinION | Illumina / 454 | ON-MinION | Sanger |
| Coverage | 4x | 174x | 80x | 107x | 50x | 80x | 55x | 131x | N/A |
| # Contigs | 79,619 | 46 | 7,398 | 84 | 26,535 | 44 | 6,044 | 21 | N/A |
| # Scaffolds | 2,051 | 31 | 220 | 55 | 237 | 27 | 111 | 17 | 8 |
| Assembly size (bp) | 43065003 | 46455514 | 41206925 | 45614181 | 42691742 | 43975886 | 41858488 | 41851079 | 41027733 |
| Largest scaffold | 557675 | 7382384 | 3113975 | 6027169 | 2814965 | 8365714 | 4783357 | 11480887 | 8319966 |
| N50 (bp) | 101645 | 5398440 | 813981 | 2812411 | 1066457 | 5531649 | 1760460 | 5096353 | 6606598 |
| N75 (bp) | 37427 | 2617860 | 277145 | 1350202 | 408473 | 3948849 | 837619 | 3936184 | 4490059 |
| L50 | 124 | 4 | 12 | 5 | 13 | 4 | 7 | 3 | 3 |
| L75 | 290 | 7 | 33 | 11 | 31 | 6 | 16 | 5 | 5 |
| BUSCO | 60.90% | 98.10% | 98.10% | 98.10% | 98.00% | 98.00% | 97.70% | 98.00% | 98.20% |
| Repeat content | 1.68% | 13.23% | 3.29% | 13.23% | 2.77% | 6.21% | 3.25% | 6.26% | 11.54% |
| GC-content | 51.39 | 50.99 | 51.09 | 50.65 | 50.96 | 49.87 | 50.81 | 50.06 | 51.61 |
| % N | 22.55 | 0.00 | 5.45 | 0.00 | 6.59 | 0.00 | 4.96 | 0.00 | 0.19 |
| # N's per 100 kbp | 22547.58 | 0 | 5445.7 | 0 | 6589.36 | 0 | 4962.16 | 0 | 189.63 |

\* from Chiapello *et al.*, 2015 [48]

### The rice, foxtail millet and goosegrass isolates of *M. oryzae* contain unique sets of mini-chromosomes

To assess the genomes for karyotype variation, we separated and visualized intact chromosomes of each strain by contour-clamped homogenous electric field (CHEF) gel electrophoresis. Strikingly, we found large-scale, structural variation in form of chromosome length polymorphisms and supernumerary mini-chromosomes in the isolates FR13, US71, and CD156 ranging in size between approximately 800 kb and 1.5 Mb (Fig 1). We hypothesized that these mini-chromosomes contribute to structural variation and might be similar to supernumerary, lineage specific chromosomes reported in other plant pathogenic fungi. It has been shown that supernumerary chromosomes can have important functions in pathogenicity, but their sequence composition and intraspecies variation in *M. oryzae* is still poorly understood.

**Fig 1.**
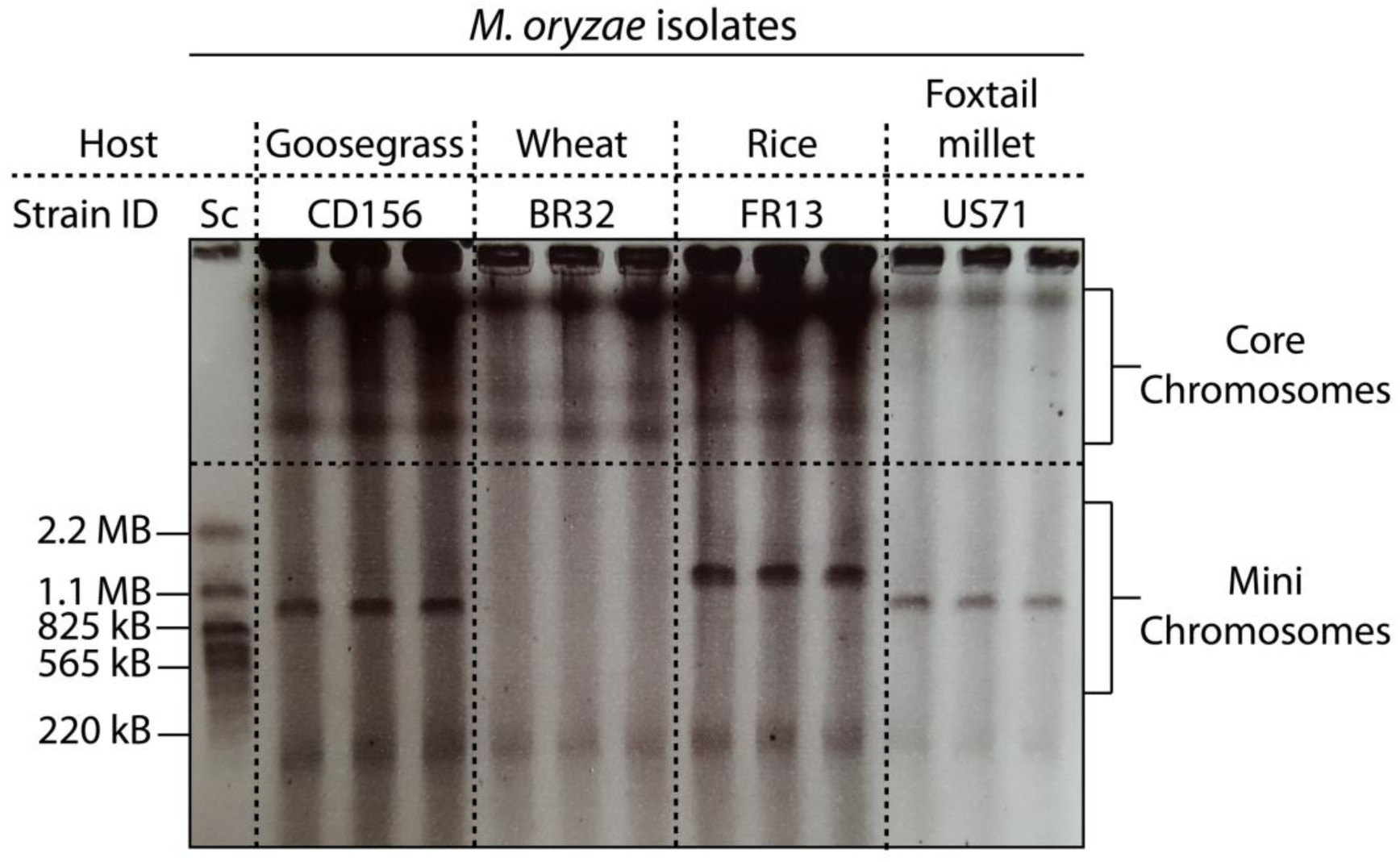
Host-specific isolates of *M. oryzae* contain mini-chromosomes of various sizes. Contour-clamped homogenous electric field (CHEF) gel electrophoresis of intact *M. oryzae* chromosomes. Chromosome size variation between *M. oryzae* isolates is present in both, mini-chromosomes and core chromosomes. Strains FR13, US71, and CD156 contain mini-chromosomes ranging in size between approximately 850 kb and 1.5 Mb. Left lane: *Saccharomyces cerevisiae* chromosomes as size marker.

To further analyze the sequence composition of the observed mini-chromosomes, we identified corresponding contigs in our whole genome assemblies by isolating mini-chromosomal DNA from CHEF gels by electro-elution for mini-chromosome isolation sequencing (detailed protocol available on protocol.io [57]; adapted from [56]). We then mapped mini-chromosome derived raw reads against the whole genome assemblies to identify specific contigs with high coverage compared to the rest of the genome, indicating their mini-chromosomal origin (Fig 2; S1, S2, S3, S4 Fig). Importantly, all core-chromosome contigs (> 2 Mb) showed extremely low coverage confirming the robustness of this approach (S1, S2, S3, S4 Fig).

**Fig 2.**
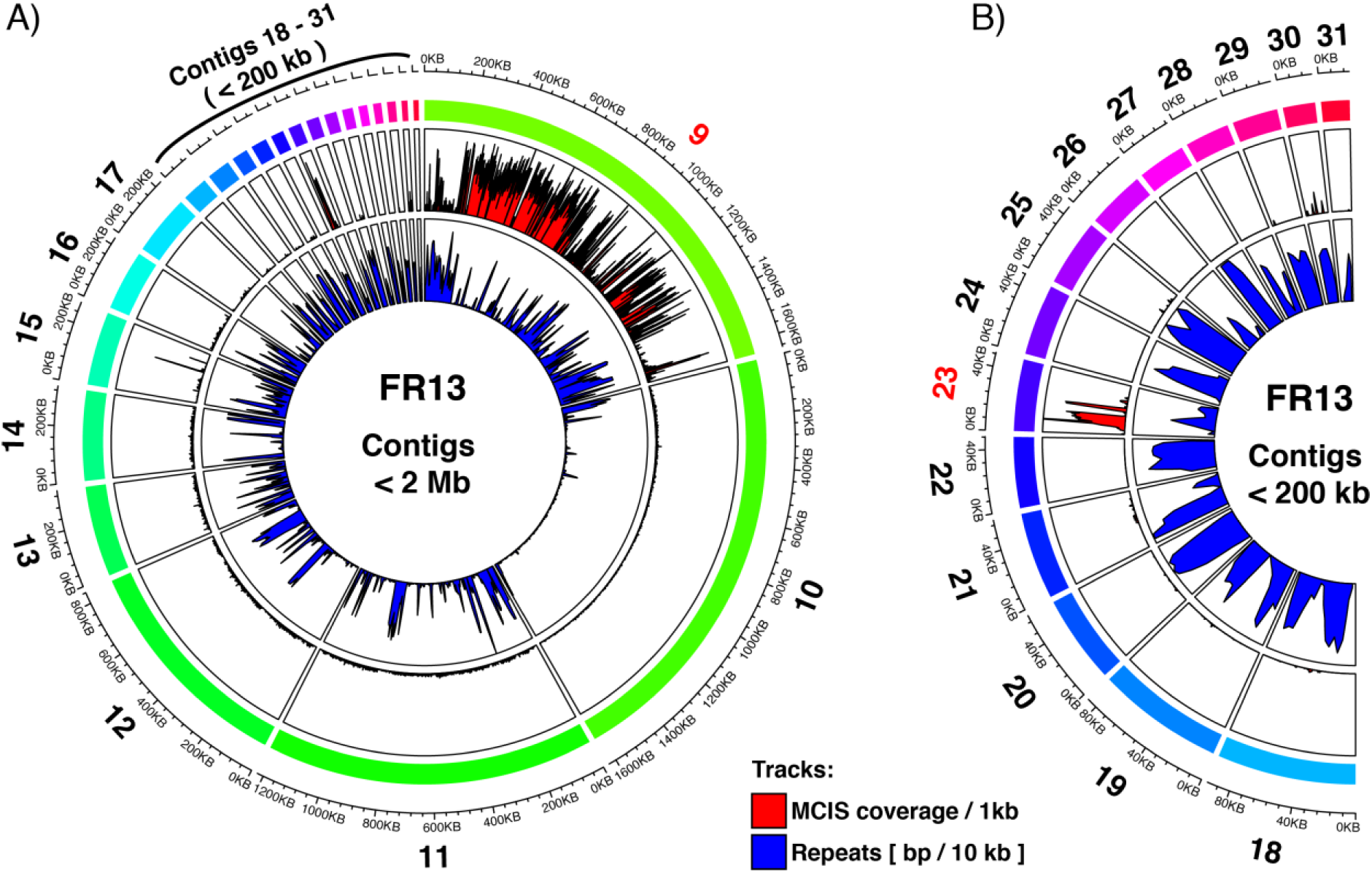
*M. oryzae* strain FR13 contains a 1.7 Mb mini-chromosome assembled into 2 contigs. **A)** Circos plot of mini-chromosome isolation sequencing (MCIS) read coverage and repeat content across FR13 contigs < 2 Mb. Outer ring (rainbow colors): FR13 contigs and contig sizes. Outer track (Red/Black): MCIS coverage per sliding window. Window size = 1000 bp; Slide distance: 500 bp. Y-axes: average coverage per 1 kb window; axis limits set to zero to maximum coverage. Inner track (Blue/Black): Repeat content per sliding window. Window size = 10 kbp; Slide distance: 5 kbp. Y-axes: repeat content in bp per 10 kb window; axis limits set to zero to maximum. B) Circos plot of MCIS read coverage and repeat content for contigs <200 kb (enlarged from A).

Repetitive sequences could lead to ambiguous mapping of reads due to the presence of similar repetitive regions in core-chromosomes. To confirm that the observed increase in coverage is linked to mini-chromosome enrichment and not to ambiguous read mapping we plotted the repeat content per 10 kb sliding window and compared it to the coverage of uniquely mapping reads derived from mini-chromosome isolation sequencing. This analysis showed that repeat-rich regions did not overlap with regions of high coverage, which confirmed that the enrichment is valid and not due to repetitive sequences and ambiguous read mapping. Using this method, we identified 2, 4, and 3 contigs with high depth and breadth of coverage for strains FR13 (Fig 2; S1 Fig), US71 (S2, S3 Fig), and CD156 (S4 Fig), respectively. The combined size of mini-chromosome contigs was 1.7 Mb for FR13, 1.58 Mb for US71, and 860 kb for CD156. For FR13 and CD156 the combined length of mini-chromosome contigs matched the size observed on CHEF gels. However, the combined length of US71 mini-chromosome contigs adds up to approximately double the size of the mini-chromosome identified by CHEF analysis, indicating that US71 most likely contains two mini-chromosomes of similar size that cannot be separated by CHEF gel electrophoresis. We also noticed an enrichment of mapped reads to the start of contig 2 in strain CD156 (S4 Fig), which might indicate that a segmental duplication contributed to the emergence of the mini-chromosome in this strain. Taken together we identified mini-chromosomes of varying size in three strains that likely represent strain-specific, genomic compartments in *M. oryzae*.

### The three examined mini-chromosomes of *M. oryzae* have different sequence composition

Based on the variation in mini-chromosome size and number we hypothesized that they represent strain-specific genome compartments with unique sequence composition. We first asked the question whether the content of mini-chromosomes is conserved in the core genome of reference strain 70-15, by globally aligning the mini-chromosomes to 70-15. However, only a 761 kb fragment of the FR13 mini-chromosome generated an alignment matching a ∼900 kb region of core chromosome 2 of 70-15, whereas mini-chromosomes of US71 and CD156 did not generate significant alignments with the reference genome.

As synteny breaks might disrupt the global alignments between the mini-chromosomes and the 70-15 genome we further mapped mini-chromosome derived raw reads against the 70-15 assembly. The ∼900 kb region on chromosome 2 of strain 70-15 showed high coverage after mapping the mini-chromosome reads of strain FR13, confirming that this region of the mini-chromosome corresponds to the end of core-chromosome 2 in 70-15. However, we did not observe unique regions with high coverage after mapping of mini-chromosome derived reads from the more distantly related strains US71 and CD156 (S5 Fig). This indicates that the mini-chromosomes of these strains contain unique sequences compared to strains FR13 and 70-15. We further examined the total amount of reads that mapped to any position in the reference genome. We found that 82.78 %, 59.77 %, and 55.49 % of mini-chromosome reads of FR13, US71, and CD156, respectively, mapped to largely overlapping regions in the reference genome, indicative for unspecific mapping. However, the fraction of unmapped reads varied between FR13 (17.21 %) and US71 (40.23 %) or CD156 (44.51 %) indicating that almost half of the mini-chromosome derived reads of US71 and CD156 originate from isolate-specific regions.

To further assess the uniqueness of mini-chromosomes between isolates we performed pairwise alignments between extracted mini-chromosome contigs of each strain. We filtered the alignments for regions that align over > 10kb to exclude unspecific, short alignments generated by repetitive regions or local similarities. Only a small fraction of the mini-chromosomes formed alignments under these parameters, whereas core-chromosomes were highly similar to each other (Fig 3A). Between CD156 and US71 only 28.39 % and 15.39 % of the mini-chromosomes generated alignments, respectively. Between FR13 and US71 the fraction of the mini-chromosomes that generate alignments was even lower with 15.01 % and 14.02 % and between FR13 and CD156 only 5.14 % or 2.61 %, respectively (Fig 3B). In contrast, between 73.88 % and 81.26 % of the core-genome generated alignments. The difference in sequence similarity between core- and mini-chromosomes supports the hypothesis that mini-chromosomes represent isolate-specific genomic entities, possibly involved in host specialization and suggests that mini-chromosomes emerged independently in host-adapted lineages.

**Fig 3.**
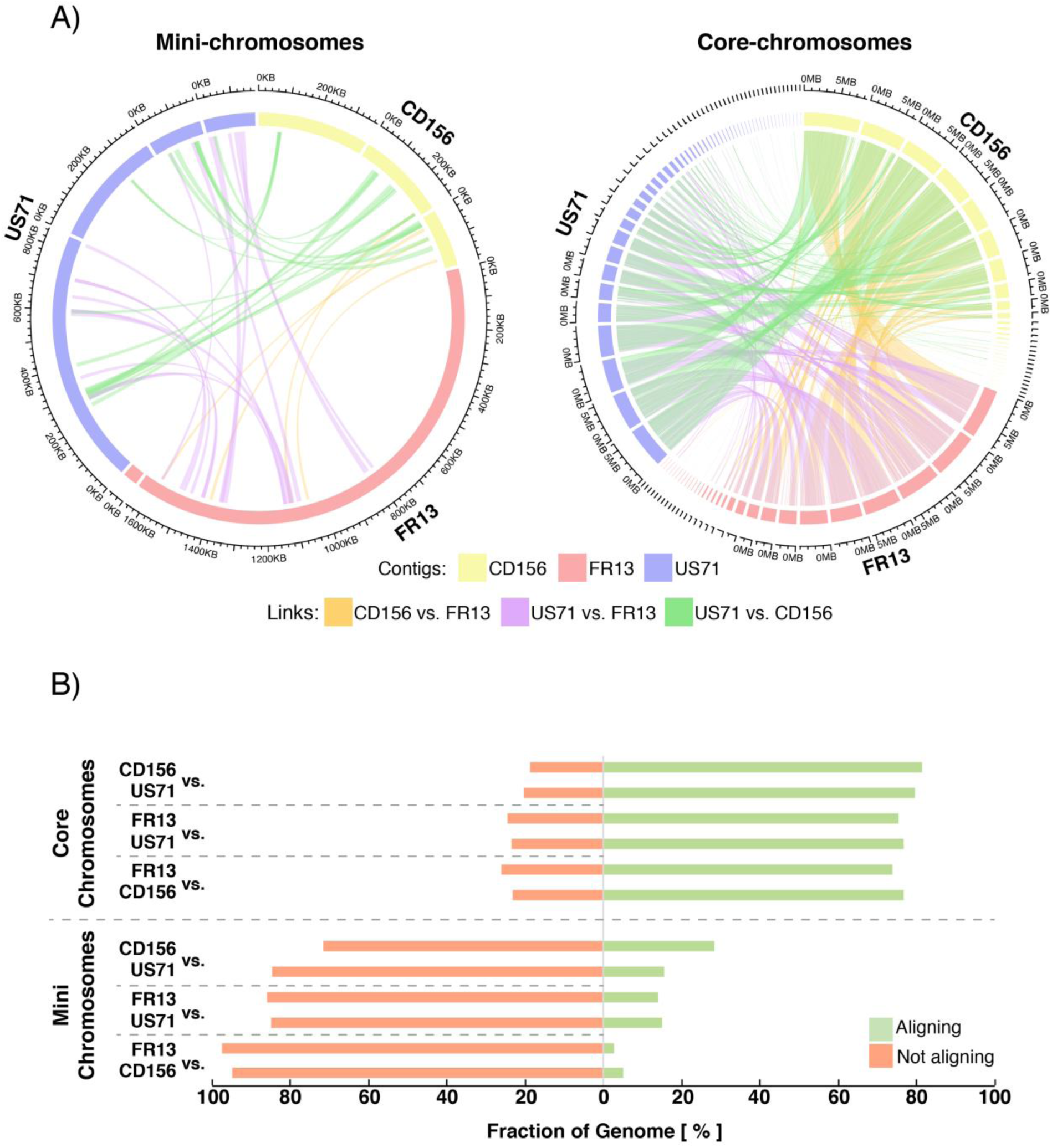
Mini-chromosomes of *M. oryzae* are isolate specific. **A)** Circos plots depicting alignments between mini-chromosomes (left) and core-chromosomes (right). Outer ring: Mini-chromosome contigs and sizes. Alignments > 10 kb are plotted as genetic links in the center. **B)** Relative fraction of mini- and core-chromosomes that generate pairwise alignments. Relative fraction of the genomic compartment that form alignments > 10 kb are shown for each pairwise alignment and each individual isolate. Approximately 75% of the core-chromosomes generate alignments under these parameters, whereas only 2.61 – 28.39% of mini-chromosomes do.

### Mini-chromosomes have lower gene and higher repeat density than core-chromosomes

To determine the gene content of the mini chromosomes, we mapped the GEMO gene annotations [48] to our new genome assemblies using BLASTn in combination with a sequence similarity approach (see methods). Of 14515, 14013 and 14415 genes that were used as queries, 13828, 13746 and 14201 mapped to a single site in the nanopore assemblies for strains FR13, US71 and CD156, respectively. Another 175, 179 and 44 genes mapped to multiple sites reflecting duplicated genes that were likely collapsed in the previous short-read assemblies. Taking these expansions into account, we assigned 14322, 14348 and 14304 genes to FR13, US71 and CD156, respectively.

Of these genes, 13963, 13985 and 14143 mapped to the core chromosomes and 359, 363 and 161 mapped to the mini chromosomes of each isolate (S1 Table). On average, in each of the three mini-chromosomes the gene content was approximately 30% lower when compared to the core genome (Fig. 4A, S1 Table).

**Fig 4.**
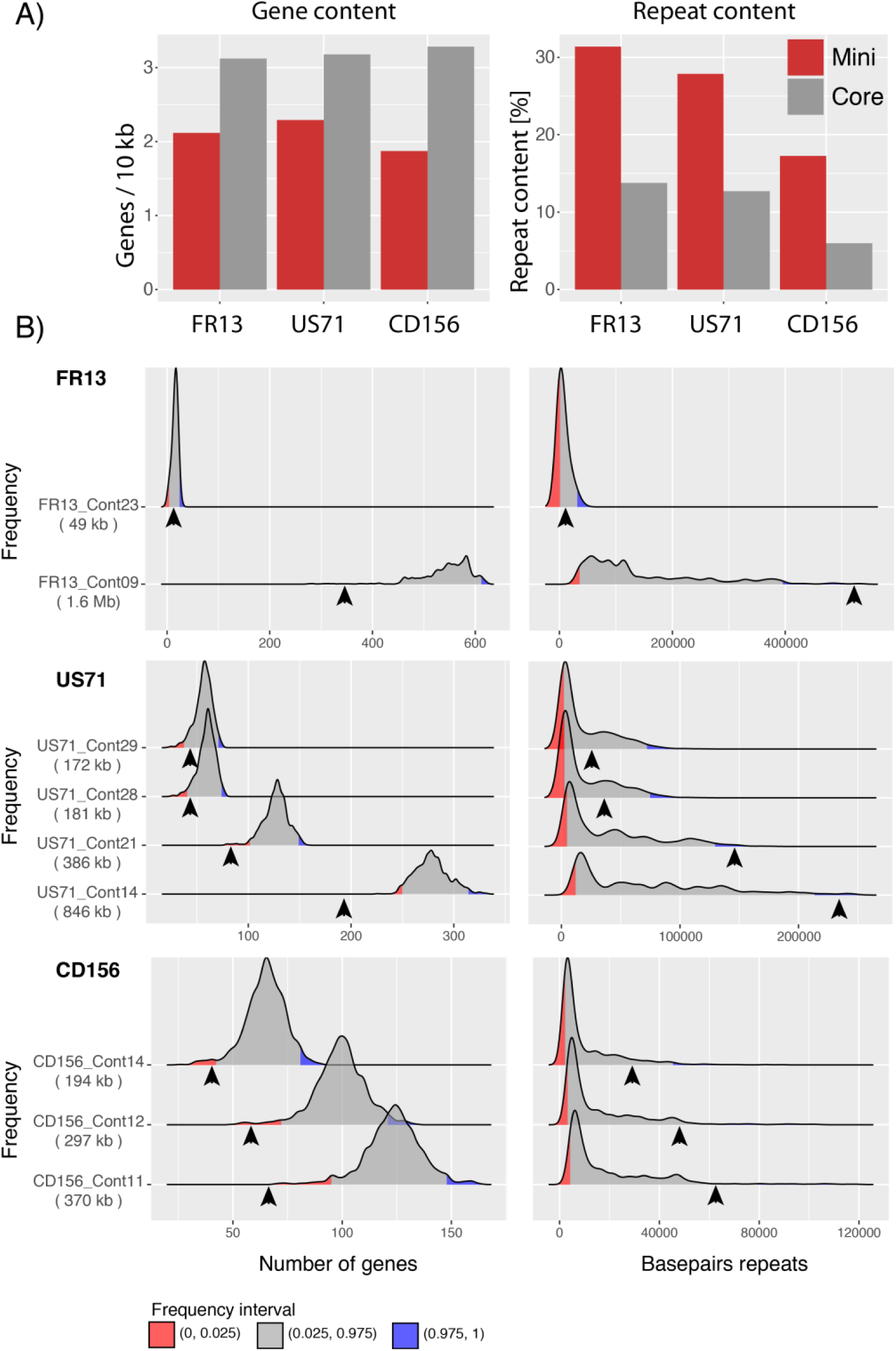
Mini-chromosomes have lower gene and higher repeat content than core-chromosomes. **A)** Bar plots of gene and repeat content of mini- (red) and core-chromosomes (grey) of isolates FR13, US71, and CD156. **B)** Density plots resulting from a bootstrapping analysis of gene and repeat content of isolates FR13, US71, and CD156. 10000 core-genomic fragments were randomly sampled for each mini-chromosome contig and distribution of genes and basepairs of repeats are shown as densities. Number of genes and repeat content of mini-chromosome contigs are indicated by arrowheads. The size of mini-chromosome contigs is given in parentheses. Lower and upper 2.5% frequency intervals are shown in red and blue. X-axis: number of genes and basepairs of repeats. Y-axis: Frequency in 10000 fragments. Y-axis limits set to min/max.

We next analyzed the repeat and GC content of the mini chromosome contigs compared to the core genomes. The GC content of core- and mini-chromosomes was similar at ∼50%, whereas the repeat content of the three mini-chromosomes differed from the core genome (Fig 4A; S1 Table). Indeed, the repeat content was more than twice as high in mini-chromosomes (31.38%, 27.87%, and 17.25%) compared to core chromosomes (13.78%, 12.71% and 5.99% in strains FR13, US71, and CD156 respectively (Fig 4A).

To exclude the possibility that the observed differences are due to biases in sample size, we performed a bootstrapping analysis for each mini-chromosome contig (see methods). Briefly, we sampled 10,000 random fragments of the size of each mini-chromosome contig from the core genome and analyzed gene and repeat content. This bootstrapping approach confirmed higher repeat and lower gene content of mini-chromosomes (Fig 4B).

These results indicate that even though the mini chromosomes have distinct sequences, they have common genomic features, low gene and high repeat density, that deviate from the typical composition of core chromosomes.

### Mini-chromosomes contain highly dynamic effector and virulence-related loci

Our gene content analysis resulted in a total of 42,974 genes between all three isolates. We grouped these genes into 12,900 orthogroups, of which 9,003 were conserved across all three strains and 3,897 were absent in at least one strain (S2 Table). Among the 3,897 orthogroups that were absent in at least one strain, we found 1,827 that were conserved between two strains and 2,070 were unique to single strains. Based on our mini-chromosome analysis we further analyzed the location of these orthogroups and categorized them into “core-genome specific”, “mini-chromosome specific” and orthogroups that contain members that are present on both core- and mini-chromosomes. We found that the vast majority (99.98%) of conserved tribes (orthogroups) was encoded exclusively on the core chromosomes or contained members on both core- and mini-chromosomes and only 0.02% are encoded exclusively on mini-chromosomes. Conversely, strain-specific tribes were slightly more abundant on mini-chromosomes (S2 Table), substantiating the observation that mini-chromosomes are strain specific and likely emerged independently in different strains or host-adapted lineages.

To determine functional categories of mini-chromosome encoded genes we performed a Hidden Markov Model (HMM) scan against the Pfam database (https://pfam.xfam.org/) using a total of 772 non-redundant, mini-chromosome encoded proteins. More than 50% of these proteins (415) did not contain any known domain, whereas 357 proteins contained Pfam domains. Certain domains were shared between at least two mini-chromosomes and included transcription factor DNA-binding domains, especially Zn(2)-Cys(6) zinc-finger, DDE-superfamily endonuclease, Tc5 transposase DNA-binding, and Methyl-transferase domains as well as domains of unknown function, amino-acid permease, protein kinase, and glycosyl-hydrolase domains. However, the relative abundance of these domains varied between individual isolates (S3 Table) and it is unclear if the presence of these protein domains on the mini-chromosomes has functional implications.

The most abundant, isolate-specific domains on mini-chromosomes were present in the rice isolate FR13 and included cytochrome P450 and polyketide-synthase domains. Both of these domains are involved in pathogenicity in *M. oryzae* and other plant pathogens [17,58]. Interestingly, we identified a well predicted polyketide synthase as Ace1 (avirulence conferring enzyme 1), which is part of a large secondary metabolite cluster and triggers an immune response in host plants carrying the resistance gene Pi33 [58].

Virulence and host adaptation of *M. oryzae* are largely determined by genes encoding secreted effector. To identify secreted proteins on mini-chromosomes, we predicted the secretome of all isolates using signalp 2.1 in combination with targetp and tmhmm to remove mitochondrial and transmembrane proteins (see Methods). This resulted in 1,394, 1,558, and 1,611 secreted proteins in strains FR13, US71, and CD156, respectively. We found 50, 8, and 2 secreted proteins encoded on mini-chromosomes, most of which correspond to uncharacterized, hypothetical proteins (S4 Table).

We further investigated whether candidate effector genes are present on the mini- and core-chromosomes using 27 characterized effectors and 167 predicted MAX-effectors (<u>M</u>agnaporthe <u>A</u>VRs and To<u>x</u>B) identified by de Guillen *et al*. [59]. MAX-effectors represent a unique class of proteins that is expanded in *M. oryzae* and contain proteins that are sequence unrelated but share a similar core structural fold. Using tblastn we identified 75, 67, and 63 proteins with similarity to known or predicted MAX-effectors in the genomes of FR13, US71, or CD156, respectively. Most of these genes were located on the core-chromosomes (S6 Fig). However, we found 5 genes in FR13 and 2 genes in US71 that were located on the mini-chromosomes. Interestingly, 3 of these genes were duplicated either on the same mini-chromosome or between the mini- and core chromosomes, suggesting structural, genomic rearrangements following segmental duplication events. Among the duplicated effector genes, we found the rice isolate specific effector AVR-Pik on the mini-chromosome of FR13 (S6A Fig). Interestingly, we found two genes encoding different variants, AVR-PikD and AVR-PikA, located towards both ends of the FR13 mini-chromosome (S6A Fig). This is consistent with previous observations by Kusaba *et al.* [55], who identified the same effector variants on a 1.6 Mb mini-chromosome in the Japanese isolate 84R-62B and might indicate that certain mini-chromosomes are maintained in *M. oryzae* populations.

### Patterns of genomic rearrangements around effector and virulence-related loci in mini-chromosomes

Our Pfam and effector analyses suggested that virulence related genes are located on mini-chromosomes and that these regions undergo rearrangements that possibly involve inter-chromosomal translocation events. To explore this, we extracted the AVR-Pik and Ace1 loci from other highly contiguous genome assemblies of the related strains 70-15, FJ81278, and Guy11 and analyzed the macro-synteny of corresponding regions for signs of genomic rearrangements. AVR-Pik is present in the reference strain 70-15 and is encoded in the sub-telomeric region at the end of chromosome 2 and on contigs 13 and 21 in strains Guy11 and FJ81278, respectively. In strain 70-15 the AVR-Pik gene resides in the region that is syntenic to the 761 kb region on the mini-chromosome of FR13 described earlier. However, comparison of the corresponding region of Guy11 suggested large scale rearrangements and variable degrees of synteny around the AVR-Pik locus in different rice isolates (Fig 5A). Additionally, we observed that the syntenic regions that are shared between the analyzed strains encode different variants of the AVR-Pik effector. Whereas the first syntenic region of the FR13 mini-chromosome encodes the variant AVR-PikA, syntenic regions in 70-15 and Guy11 encode the variant AVR-PikC. Furthermore, we identified a 82 kb region in isolate FJ81278 encoding AVR-PikD with high sequence similarity and synteny to the end of the FR13 mini-chromosome that encodes the same AVR-Pik variant, suggesting that the two copies of AVR-Pik on the mini-chromosome originated from independent genomic locations and recombined on the mini-chromosome.

**Fig 5.**
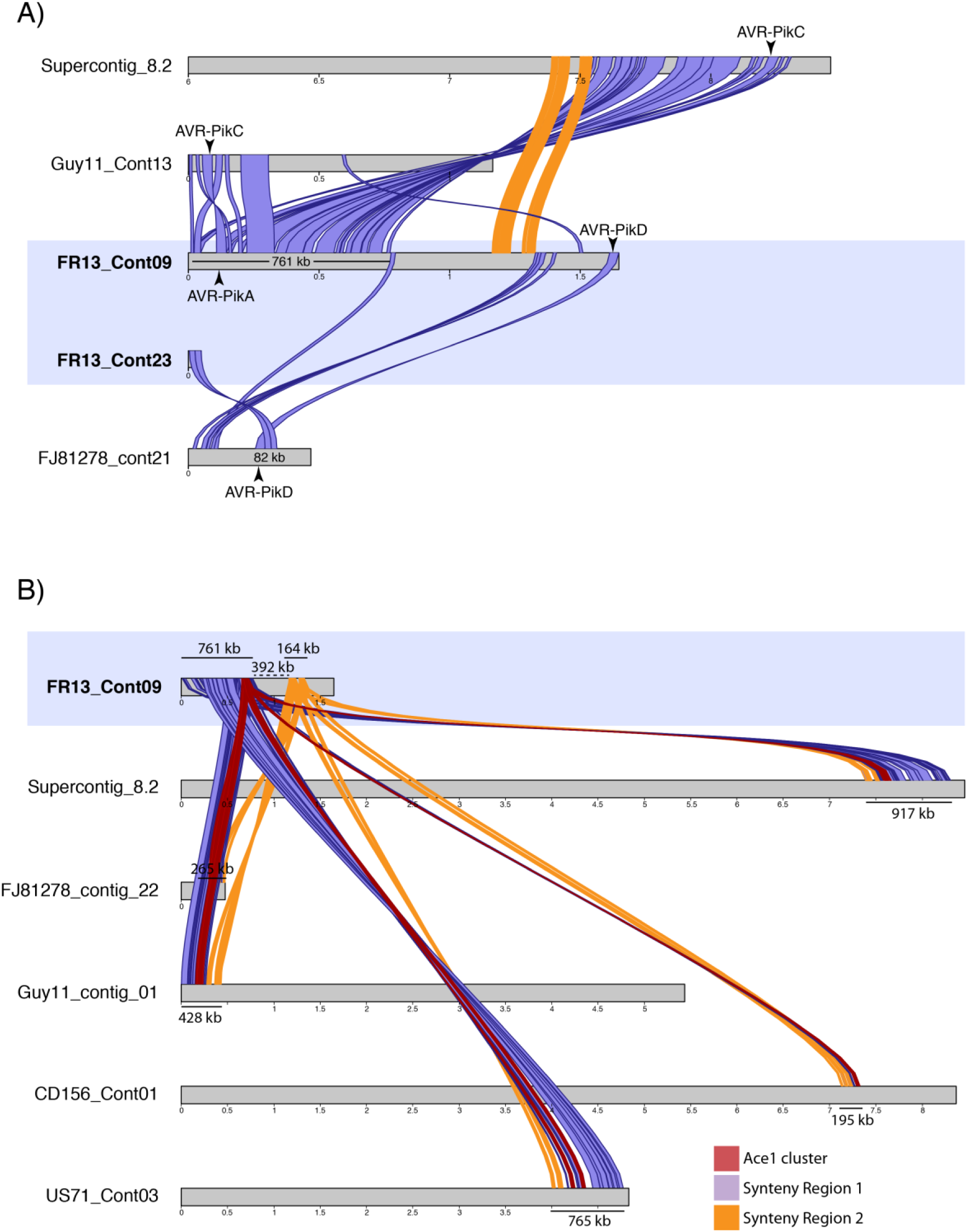
Genomic rearrangements around virulence-related loci on mini-chromosomes. **A)** Synteny analysis around the AVR-Pik loci on the FR13 mini-chromosome and corresponding chromosomes and contigs in the related isolates Guy11, FJ81278, and the reference strain 70-15. The 761 kb region around the variant AVR-PikA on the mini-chromosome with high similarity to the end of chromosome 2 of the reference strain 70-15 is shown in purple. The makrosyntenic region between 70-15 and FR13 is disrupted on the mini-chromosome (synteny break between region 1 and 2). The synteny is disrupted earlier in Guy11 and there are 2 inversions around the AVR-Pik locus. Another syntenic region (82 kb) is present between the end of the mini-chromosome encoding the AVR-PikD variant and contig 21 of isolate FJ81278. Syntenic regions larger than 10 kb are shown. Illustration of supercontig 8.2 starts at 6 Mb for better visualization. **B)** Synteny analysis around the Ace1-containing secondary metabolite cluster on the FR13 mini-chromosome and corresponding contigs of isolates Guy11, FJ81278, 70-15, US71, and CD156. The Ace1 cluster is located at the end of the 761 kb syntenic region between the FR13 mini-chromosome and supercontig 8.2 of isolate 70-15. The synteny around the Ace1 locus is highly conserved in closely related isolates, including the foxtail millet isolate US71. In strains Guy11, US71, and 70-15 the cluster is located on core-chromosomes. The macro-synteny around the Ace1 cluster is disrupted by a 392 kb insertion on the FR13 mini-chromosome. The Ace1 cluster is shown in red and the two syntenic regions (761 kb and 164 kb) are shown in purple and yellow, respectively. Mini-chromosome contigs are highlighted in blue.

The polyketide synthase Ace1 is also located on the FR13 mini-chromosome. Ace1 is part of a large secondary metabolite cluster that spans a region of approximately 70 kb [58]. To analyze the genomic context around this cluster on the mini-chromosome, we extracted the gene sequences of the whole cluster in FR13 and analyzed syntenic regions in the aforementioned isolates as well as US71 and CD156. In rice isolates and the foxtail millet isolate US71 the Ace1 cluster and the order of the macrosyntenic region are well conserved (Fig 5B). Interestingly, the ACE1 cluster is encoded on core-chromosomes in strains 70-15, Guy11, and US71, based on the size of the contigs, suggesting inter-chromosomal rearrangements that involve core- and mini-chromosomes. In the more distantly related strain CD156 the Ace1 cluster is less conserved and the syntenic region only spans across 195 kb, reflecting the genetic diversity between rice isolates and distantly related lineages. Strikingly, the macrosyntenic region on the FR13 mini-chromosome is disrupted by a 392 kb insertion after the Ace1 cluster that is absent in the core-chromosomes of other isolates, further substantiating the hypothesis that mini-chromosomes might facilitate genomic rearrangements.

## Discussion

Genomes of eukaryotic plant pathogens are notorious for being highly dynamic. Chromosome scale, structural variation, including supernumerary mini-chromosomes, has been documented by electrophoretic karyotyping [14] but has not been studied extensively at the sequence level, mainly due to methodological limitations. Here, we took advantage of recent technical developments to reassess the genomes of 4 previously sequenced, host-specific isolates of the blast fungus *M. oryzae*. We used a combination of long- and short-read sequence data to generate near chromosome quality genome assemblies. We found that supernumerary mini-chromosomes are common in the independent lineages of the blast fungus *M. oryzae*. We then used a technique we termed mini-chromosome isolation sequencing to collect short-read data from isolated mini-chromosomes and match them to contigs in the whole genome assemblies. We detected single mini-chromosomes in the rice and goosegrass isolates and two mini-chromosomes in the foxtail millet isolate of *M. oryzae* and discovered that the sequence of the mini-chromosomes is highly variable between strains, suggesting that they emerged independently probably from rearrangements of core-chromosomes. However, the mini-chromosomes share common features such as low gene and high repeat density. Additionally, we located the genes for the effector AVR-Pik and the polyketide synthase Ace1 to the mini-chromosome of the rice-infecting isolate FR13 even though these genes are located on core-chromosomes in other isolates. Analyses of macrosynteny around these regions showed that mini-chromosomes undergo large-scale rearrangements, and thus, could contribute to genomic plasticity within *M. oryzae* populations. Overall, our results demonstrate the value of mini-chromosome isolation sequencing as a scalable method to reliably identify mini-chromosomes in whole genome assemblies. This will allow us to study this unique genomic compartment at the sequence level and help to unravel the mechanism that compartmentalizes plant pathogen genomes.

Our study, along with Peng et al. [56], reveals that mini-chromosomes are present in independent lineages of *M. oryzae*. Although we found mini-chromosomes in the examined rice, foxtail millet and goosegrass isolates of *M. oryzae*, we didn’t detect any in the wheat blast isolate BR32 (Fig 1), an isolate that traces back to early outbreaks of wheat blast in 1991. This is consistent with the recent study by Peng *et al*. [56], which documented mini-chromosomes in the wheat blast isolates B71 and P3 collected in 2012 but not in T25 collected in 1988. It is possible that wheat blast isolates collected soon after the emergence of this disease in the 1980s lack mini-chromosomes. The lack of meiotically unstable mini-chromosomes in these isolates may be due to sexual reproduction, which was common among isolates from the early phases of the epidemic [60]. However, we cannot rule out the possibility that these isolates have lost their mini-chromosomes over time due to long term culturing under laboratory conditions. Similar loss of accessory chromosomes within weeks has been demonstrated *in vitro* for *Z. tritici* [20]. The observed variation in mini-chromosome content is consistent with earlier studies showing that the occurrence of mini-chromosomes is variable within and between lineages of *M. oryzae* [52,53].

The three mini-chromosomes we studied share little sequence similarity with each other indicating that they are not conserved across host specific lineages but rather emerged independently throughout the diversification of *M. oryzae* (Fig 3). In contrast, mapping of mini-chromosome derived reads of wheat isolate P3 to the B71 mini-chromosome sequence revealed several overlapping regions [56]. Despite the overlaps, these regions seem to be interspersed with sequences that are specific to each individual mini-chromosome. It is likely that mini-chromosome divergence follows overall genetic distance between isolates explaining the low sequence similarity between the mini-chromosomes we sequenced. This would be consistent with the view that mini-chromosomes emerge from rearrangements of core-chromosomes. However, given that vast variation in mini-chromosome size was observed among rice isolates [52], variability of mini-chromosomes within host specific lineages is likely to occur. Future comparative analyses of natural populations of *M. oryzae* will shed light on the mechanisms of mini-chromosome evolution.

Despite their independent origin, mini-chromosomes share common genetic features. We notice that elevated repeat content and lower gene densities are consistent features of mini-chromosomes (Fig 4) [56]. This is surprising given that all mini-chromosomes share overall little sequence similarity and seem to have emerged independently. What is the underlying mechanism that facilitates emergence of sequence unrelated mini-chromosomes with common genomic features? Our results suggest that mini-chromosomes might emerge from core genomic rearrangements involving primarily core-chromosome ends (Fig 5; S4 Fig). These regions are known to contain higher repeat content and to be more dynamic than central regions of chromosomes [11,56,61]. It seems plausible that repeats are involved in mini-chromosome formation. Repeats facilitate genomic rearrangements, such as inter-chromosomal translocations, in several plant pathogenic fungi [6,11,20,62,63]. Under stress conditions, specific classes of transposable elements can get activated and induce genomic rearrangements [64]. Interestingly, Peng et al. [56] observed that the B71 mini-chromosome contains more active repeats than core chromosomes based on lower rates of repeat-induced point (RIP) mutations. The emerging model is that transposon activation generates genomic instability, primarily at core-chromosome ends, resulting in the genesis of mini-chromosomes. Future comparative analyses of mini-chromosomes from closely related isolates, ideally within the same clonal lineage, will further reveal the precise genetic elements associated with the emergence and divergence of mini-chromosomes.

What are the implications of mini-chromosomes for adaptive evolution of the blast fungus? Our results, together with other’s suggest that mini-chromosomes form a gene poor, repeat-rich genomic compartment that contributes to genome rearrangements and likely gene flow [11,55,56]. Given that effector genes are associated with mini-chromosomes of *M. oryzae*, this genomic architecture is another example of the two-speed genome concept in which particular compartments contribute to adaptive evolution. This could happen through several mechanisms. Mini-chromosomes may represent an intermediate stage of large rearrangements that facilitate generation of structural variations across the genome. Additionally, mini-chromosomes might facilitate transfer of genetic material between individuals by enabling gene flow independently of core genomic recombination. Transfer of supernumerary chromosomes has been reported for several plant pathogenic fungi [19,27,55,65]. A species such as *M. oryzae* that consists of genetically defined host-specific lineages may benefit from a process that facilitates horizontal transfer of genetic material. Genetically diverse isolates that infect a common host may give rise to new variants that increase the diversity and adaptive potential of local blast populations [66]. Therefore, it is of utter importance to understand the biology of supernumerary mini-chromosomes and its impact on genomic diversity and gene flow. Our study lays the basis for studying the role of mini-chromosomes in defining the genetic identity of host specific lineages of *M. oryzae*. Future analyses of a larger number of field isolates will further define the degree to which mini-chromosomes shape the evolution of host-specific lineages of *M. oryzae*.

## Material and Methods

### Biological material

The *M. oryzae* isolates analyzed in this study were field isolates collected from different hosts and different regions and represent four host-adapted lineages. FR13 was isolated from japonica rice in France in 1988, US71 was isolated from *Setaria sp*. (foxtail millet) in the USA, CD156 was isolated from *Eleusine indica* (goosegrass) in Ferkessedougou, Ivory Coast in 1989, and BR32 was isolated from wheat in Brazil in 1991. All isolates were acquired from Elisabeth Fournier and have been previously reported in the GEMO project [48].

### DNA extraction for whole genome sequencing and mini-chromosome isolation sequencing

For mycelium propagation and whole genome sequencing, *M. oryzae* isolates were cultured on complete medium agar (CM). High molecular weight genomic DNA from *M. oryzae* was extracted from mycelia of 7-day old cultures following the method described in [67]. Genomic DNA was quantified on a TapeStation (Agilent) and treated with DNAse-free RNAse. RNAse treated DNA was sheared using either a gTUBE or a 22 Gauge needle. Sheared DNA was captured using AMPure beads (Beckman Coulter, Indianapolis, US) and eluted in 45 μl water.

For mini-chromosome isolation sequencing, 8 blocks of 7 days old mycelium were transferred into 150 ml YG-medium (5g/L yeast extract, 20g/L glucose) and incubated at 24°C and 120 rpm for 3 days. Mycelium was harvested by filtering the culture through two layers of miracloth. Protoplasts were generated by incubation in sterile *Trichoderma harzianum* lysing enzymes solution (150 mg lysing enzymes in 15 mL 1 M Sorbitol) for 2-4 h. Protoplasts were harvested by filtering through two layers of miracloth followed by centrifugation for 10 min at 1,500 rpm and washed twice in 1 M Sorbitol. Quality of protoplasts was observed microscopically. Protoplasts were then resuspended in 100-200 µl 1 M Sorbitol / 50 mM EDTA, embedded in 2x volumes of 1% certified megabase agarose (Biorad) and incubated in proteinase K containing NDS buffer (10 mg/mL laurylsarcosine, 100 mM TRIS-HCl pH9.5, 500 mM EDTA, proteinase K 200 µg/ml) at 50°C for 48 h. Plugs were then washed three times with fresh 50 mM EDTA for 1 h prior to CHEF gel electrophoresis (detailed protocol available on protols.io, [68]).

### Whole genome sequencing and assembly

Libraries for whole genome sequencing were prepared by following the 1D protocol from Oxford Nanopore. Sequencing runs were performed using MinION R9.4 (Oxford Nanopore Technologies, Oxford, UK). Sequence reads were assembled into contigs using Canu (v1.6 and v1.7) [69]. Contiguity was further improved by merging highly similar contig ends with identity >98% and alignment length >10 kb (for three contigs the length of the alignment was between 8 kb and 10 kb) after whole genome alignments using the nucmer application of the MUMMER3 package [70] and SSPACE [71]. After extracting and merging matching contig ends we tested for read support for both ends and reads spanning the entire region (Sup. Table 5). We then improved base calling quality of the assemblies by applying a polishing pipeline (https://github.com/nanoporetech/ont-assembly-polish) consisting of two iterations of racon (https://github.com/isovic/racon; v1.3.2) and two iterations of pilon v1.22 [72] polishing using Nanopore and published Illumina reads (Illumina reads from [48]). To assess the performance of the polishing pipeline and the overall quality of the genome assemblies we performed a benchmarking universal single-copy orthologs (BUSCO) [73] analysis using the sordariomyceta database (https://busco.ezlab.org/). Details about the assemblies, isolates, and accession numbers are available in [57]. Nanopore sequencing reads are deposited in the European Nucleotide Archive (ENA) under the accession numbers ERR2612751 (BR32), ERR2612749 (FR13), ERR2612750 (US71), and ERR2612752 (CD156).

### Mini-chromosome isolation, library preparation and sequencing

Mini-chromosomes of *M. oryzae* were separated from core-chromosomes by contour-clamped homogenous electric field (CHEF) gel electrophoresis. Therefore, plugs containing the digested protoplasts (see above) were placed in a 1% megabase agarose gel. We used 0.5% TAE buffer for subsequent DNA extraction or 0.5% TBE buffer for visualization (Fig. 1). We separated mini-chromosomes using a Biorad CHEF DRII system with the following settings: Run time: 96 h; Voltage: 1.8-2 V/cm; Initial switch interval: 120 s; End switch interval: 3600 s. The gel was then dyed with ethidium bromide solution (1 µg/ml) for visualization on a UV-transilluminator and individual mini-chromosome bands were excised.

Mini-chromosomal DNA was eluted from the gel plugs by electroelution using a D-Tube dialyzer midi, MWCO 3.5 kD (Merck). Therefore, individual plugs were placed in the dialysis tube and covered with 0.5% TAE buffer. The mini-chromosomal DNA was electroeluted for 3 h at 90 V, resuspended by slowly pipetting up and down and concentrated in a vacuum centrifugal evaporator to a concentration of 50-150 ng/µl. Concentration of the extracted DNA was measured spectrophotometrically and fluorometrically by Nanodrop and Qubit, respectively, prior to library preparation.

Libraries for mini-chromosome sequencing were generated following the general guidelines of the Nextera Flex library preparation kit (Illumina). We modified the protocol as follows. For tagmentation we used a total volume of 5 µl consisting of 2.5 µl Tn5 transposase, 0.5 µl of 10x reaction buffer, and 2 µl mini-chromosomal DNA set to a concentration of 0.5 ng/µl. The tagmentation mix was incubated in a thermocycler at 55°C for 7 min. The tagmentation product was then used in a PCR reaction containing 2.5 µl custom barcoding primers (10 µM) (Sup. Table 6), 25 µl 2x NEBNext High-fidelity PCR mix (New England Biolabs), and 15 µl H2O. PCR conditions: 72°C for 5 min, 98°C for 30 sec, 5 cycles of 98°C for 10 sec followed by 63°C for 30 sec and 72°C for 1 min. The numbers of amplification cycles needed for library preparation of each sample was then determined by quantitative PCR as follows: 25 µl SYBR Green Jumpstart Taq ReadyMic (Sigma-Aldrich), 5µl PCR product from previous reaction, 2.5 µl of each barcoding primer, 15 µl H2O. PCR conditions as described above, without the initial 72°C step. Same PCR conditions were then applied to amplify the tagmented DNA. Libraries were then cleaned up using AMPure PB Bead purification kit (Pacific Biosciences). Concentration and quality of the libraries was confirmed by Nanodrop, Qubit, and BioAnalyzer 2100 using the high sensitivity DNA Kit (Agilent Technologies). Sequencing of mini-chromosomal DNA libraries was carried out on a NextSeq 500 system (Illumina) using the NextSeq 500/550 Mid output Kit (Illumina). Mini-chromosome derived reads were deposited at the European nucleotide archive under the accession numbers ERR3771227-ERR3771238.

### Identification of mini-chromosomes in whole genome assemblies

The quality of reads obtained from mini-chromosome isolation sequencing was confirmed with fastQC (https://www.bioinformatics.babraham.ac.uk/projects/fastqc/) and low quality sequences as well as adaptor sequences were removed using trimmomatic [74]. Mini-chromosome reads of each strain were then mapped against the whole genome assembly of the same strain using the BWA-mem (burrows-wheeler aligner) algorithm with default parameters. Reads were then filtered to keep only uniquely mapping reads using the samtools package [75] to prevent ambiguous mapping of reads originating from repeat rich regions. MCIS read coverage was calculated per 1 kb sliding window (window size: 1000 bp; slide: 500 bp) using the bedtools package (https://bedtools.readthedocs.io/en/latest/) and plotted using the R package circlize [76]. Mini-chromosome coverage and repeat content used to generate circus plots are shown in S7 Table.

Read mapping against the reference genome of strain 70-15 and visualization was carried out as described above. The number of reads mapping to the reference genome was calculated for total reads and uniquely mapping reads. Uniquely mapping reads were used for visualization to exclude ambiguous mapping sites. Total amount of mapping reads is reported as percentage of total reads per sample.

### Whole genome and mini-chromosome alignments

Global alignments were generated using the nucmer algorithm of the MUMMER3 package [70]. The alignments were further filtered using the delta-filter utility (MUMMER3) with parameters -l 10000 and -i 80 (length >10 kb; percent identity >80%) to retrieve continuous alignments. Alignment coordinates were extracted with the show-coords utility and the output was modified to generate coordinate files in “bed” format for plotting in R. Plots were generated in R using the packages circlize [76], for circular representations of mini- and core-chromosome alignments, and karyoploteR [77], for linear representation of syntenic regions.

### Analysis of gene and repeat content

Gene models for all strains were retrieved from previous assemblies published via the *Magnaporthe* GEMO database ([48]; http://genome.jouy.inra.fr/gemo/). We then identified genes using a similarity-based approach. We used blastn [78] with all strain specific gene models to identify genes in our assemblies using a threshold of >90 query coverage and 99% identity to account for differences in base calling quality and sequencing errors between two assemblies of the same strain. The >90 coverage threshold was chosen to account for errors in mononucleotide repeats that can occur from nanopore sequencing and the 99% identity threshold was empirically determined by whole genome alignments of both assemblies of the same strain which resulted in an average sequence identity between 99% and 99.5%. Resulting genes were assigned to mini- and core-chromosome encoded. Gene density was calculated as number of features per 10 kb window across each genomic compartment (core- and mini-chromosomes).

Repetitive sequences were annotated with RepeatMasker (http://www.repeatmasker.org/) using a merged library of repeats consisting of the RepBase repeat library for fungi (https://www.girinst.org/repbase/) and *Magnaporthe* repeats identified by Chiapello et al. [48]. Total repeat content for the whole genome as well as for mini- and core-chromosome contigs was extracted from RepeatMasker.

For bootstrapping of gene and repeat content we extracted the coordinates of features from the gene model blastn and RepeatMasker output. We then transformed the data into bed format for analysis in R. For each mini-chromosome contig we sampled 10,000 random regions from the core genome with equal size to the respective mini-chromosome contig using the randomizeRegions function of the regionR package. We then determined the numbers of genes and the basepairs occupied by repeats for each core-chromosome fragment and the mini-chromosome contigs using the countOverlaps function of the GenomicRanges package. Density plots of the frequency of features per core-chromosome fragment were generated with the R package ggplot.

### Prediction of secreted proteins

To predict secreted proteins, we retrieved the protein annotation of all strains from the *Magnaporthe* GEMO database ([48]; http://genome.jouy.inra.fr/gemo/). We then predicted proteins containing a signal peptide using the program SignalP v2.1 [79]. We then used TargetP-2.0 [80] to identify mitochondrial proteins and the hidden-markov model TMHMM [81] to identify proteins containing transmembrane domains. After removing mitochondrial and transmembrane proteins we matched secreted proteins to the blastn output used to transfer gene models (described above) to determine the number and genomic location of secreted proteins in the nanopore/canu assemblies.

### Pfam domain annotation

Pfam domains were predicted by a hidden-markov model scan using the Pfam protein families database ([82]; https://pfam.xfam.org/). Protein sequences were retrieved from the *Magnaporthe* GEMO database as described above.

### Predicting orthologous groups by TRIBE-MCL

Orthologous families were predicted using the software TRIBE-MCL [83]; http://micans.org/mcl/). We combined the proteomes of all strains and identified homologous proteins using blastp with a query coverage threshold of 90% and e-value of 10e^10. We then transform the blastp output into an MCL readable format using the mcxdeblast command of TRIBE-MCL and identify tribes by mcl and extracted tribes and protein identifiers using custom perl scripts. Of 12951 tribes predicted from the combined proteome retrieved from the GEMO Magnaporthe database, the members of 12900 matched the new assemblies with high coverage (>90%) and identity (>99%). Resulting tribes were then categorized into three groups: i) tribes that are conserved in all isolates, ii) tribes that are missing in one of the isolates, and iii) tribes only present in one isolate. Then we assigned the genomic location of each of the tribe members as core-chromosome or mini-chromosome encoded, based on our mini-chromosome identification.

## Acknowledgements

We thank Elisabeth Fournier for providing the isolates used in this study, Florian Charriat and Pierre Gladieux for providing the repeat library used to annotate repetitive regions, Pingtao Ding for helpful advice and support during library preparation and mini-chromosome sequencing, Dan MacLean and Clémence Marchal for bioinformatics support, and Nick Talbot, Hernan Burbano, and Angus Malmgren for critically reading the manuscript. This work was supported by the Gatsby Charitable Foundation, the ERC (proposal BLASTOFF 743165), and BBSRC.

## Author Contributions

**Conceptualization:** Sophien Kamoun, Thorsten Langner

**Data acquisition and curation:** Thorsten Langner, Adeline Harant, Luis B. Gomez-Luciano, Ram K. Shrestha, Joe Win

**Handling of biological material:** Adeline Harant, Thorsten Langner, Joe Win

**Data Analysis:** Thorsten Langner, Luis B. Gomez-Luciano, Joe Win

**Supervision:** Sophien Kamoun

**Manuscript preparation:** Thorsten Langner, Sophien Kamoun

**Manuscript review:** Thorsten Langner, Sophien Kamoun, Adeline Harant, Luis B. Gomez-Luciano, Ram K. Shrestha, Joe Win

**Funding acquisition:** Sophien Kamoun

## Data availability

All sequence data used in this study was deposited at the European Nucleotide Archive (ENA) with the study accession PRJEB27137. Nanopore sequencing reads were deposited under accession numbers ERR2612751 (BR32), ERR2612749 (FR13), ERR2612750 (US71), and ERR2612752 (CD156). Nanopore sequence assemblies were deposited under accession numbers GCA_900474545.2 (BR32), GCA_900474655.2 (FR13), GCA_900474175.2 (US71), and GCA_900474475.2 (CD156). Polished whole genome assemblies were deposited under accession numbers GCA_900474545.3 (BR32), GCA_900474655.3 (FR13), GCA_900474175.3 (US71), and GCA_900474475.3 (CD156). Mini-chromosome isolation sequencing reads were deposited under accession numbers ERR3771227-ERR3771238.

## Supplementary information

**S1 Fig.**
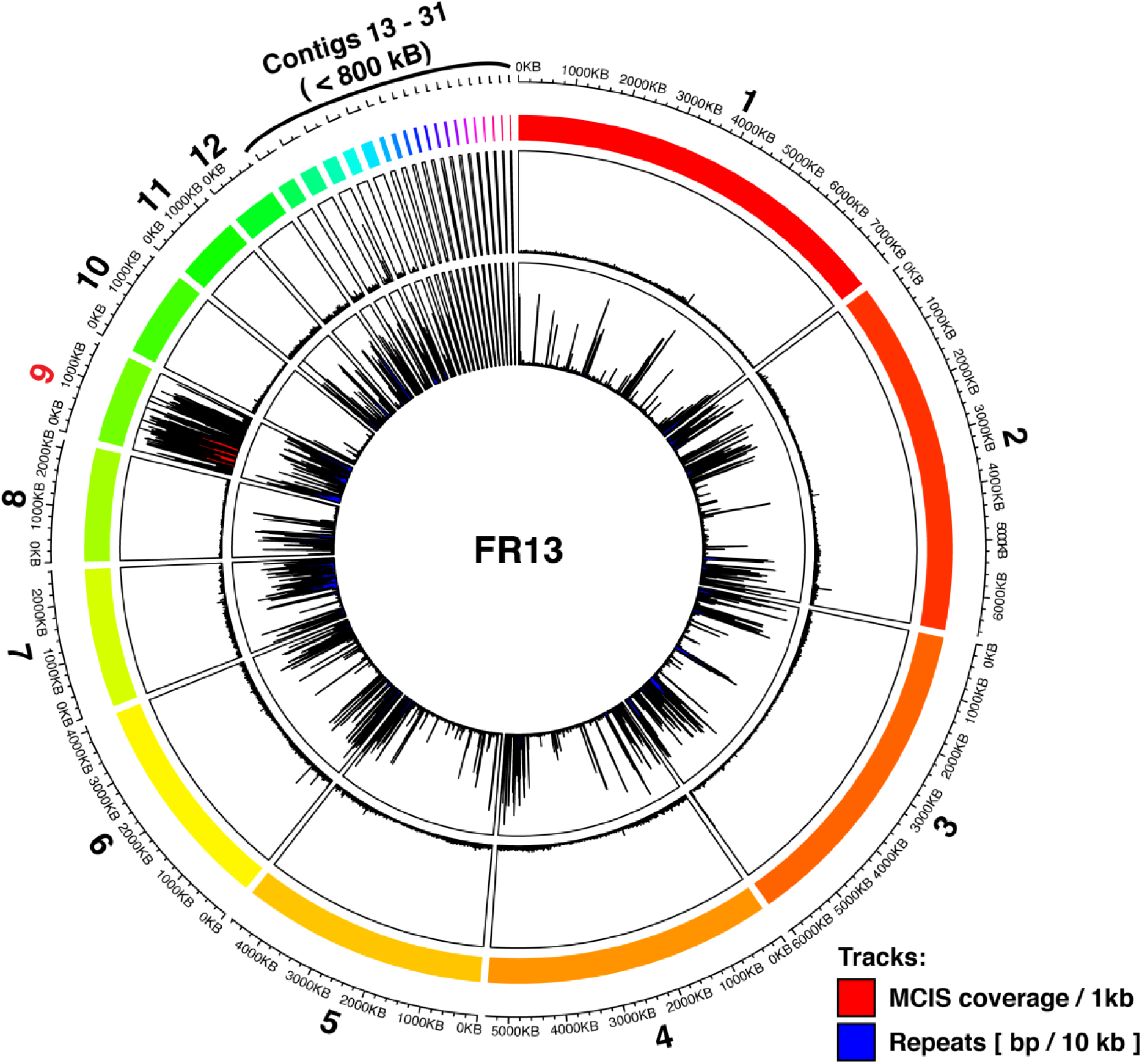
MCIS-read mapping and repeat content of *M. oryzae* strain FR13. Circos plot of mini-chromosome isolation sequencing (MCIS) coverage and repeat content across the FR13 genome. Outer ring (rainbow colors): Contigs and contig sizes. Outer track (Red/Black): MCIS coverage per sliding window. Window size = 1000 bp; Slide distance: 500 bp. Y-axes: average coverage per 1 kb window; axis limits set to min/max coverage. Inner track (Blue/Black): Repeat content per sliding window. Window size = 10 kbp; Slide distance: 5 kbp. Y-axes: repeat content in bp per 10 kb window; axis limits set to zero to maximum.

**S2 Fig.**
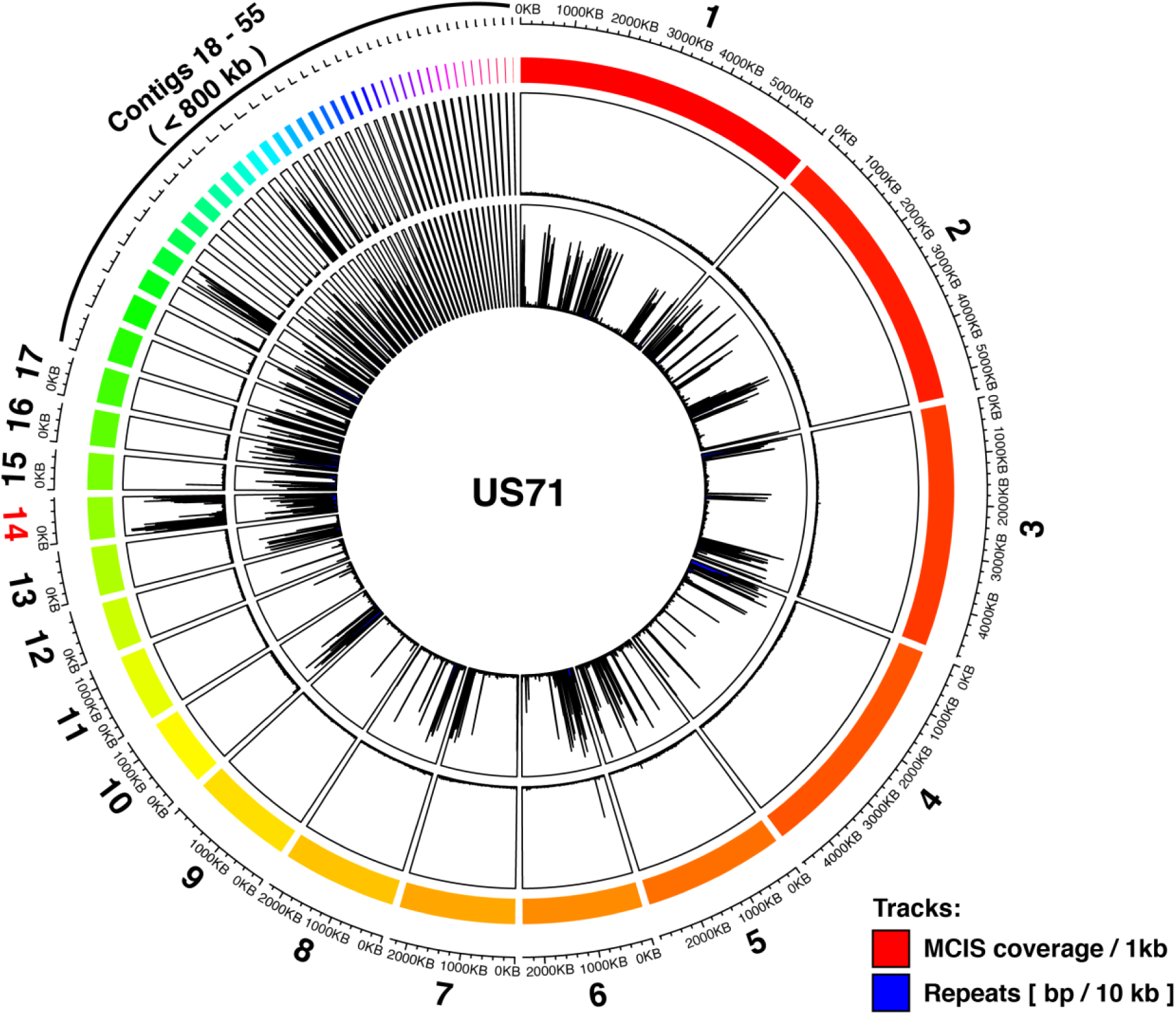
MCIS-read mapping and repeat content of *M. oryzae* strain US71. Circos plot of mini-chromosome isolation sequencing (MCIS) coverage and repeat content across the US71 genome. Outer ring (rainbow colors): Contigs and contig sizes. Outer track (Red/Black): MCIS coverage per sliding window. Window size = 1000 bp; Slide distance: 500 bp. Y-axes: average coverage per 1 kb window; axis limits set to min/max coverage. Inner track (Blue/Black): Repeat content per sliding window. Window size = 10 kbp; Slide distance: 5 kbp. Y-axes: repeat content in bp per 10 kb window; axis limits set to zero to maximum.

**S3 Fig.**
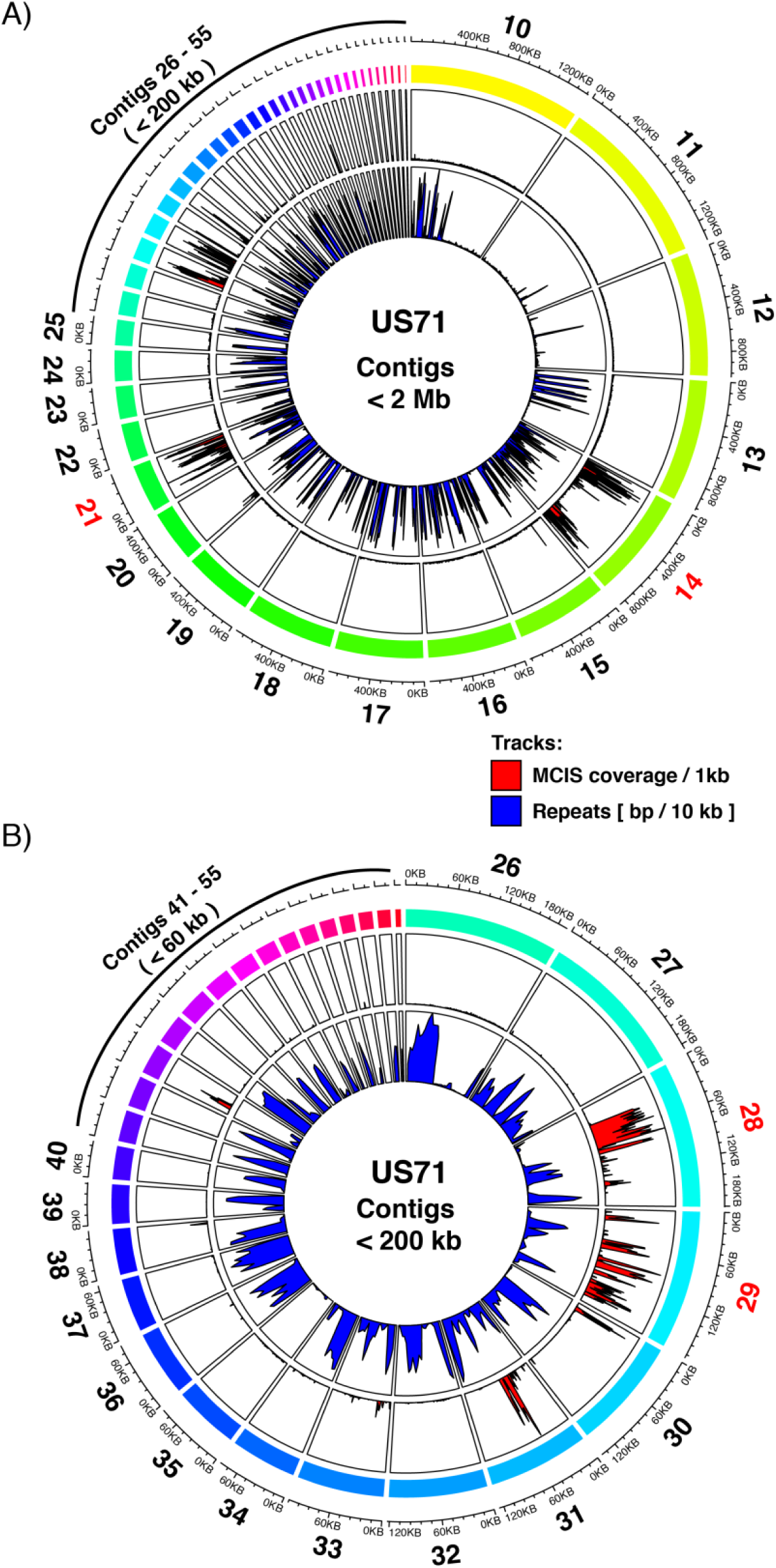
MCIS-read mapping and repeat content of contigs < 2 Mb in *M. oryzae* strain US71. **A)** Circos plot of mini-chromosome isolation sequencing (MCIS) coverage and repeat content across US71 contigs < 2Mb. **B)** Circos plot of mini-chromosome isolation sequencing (MCIS) coverage and repeat content across US71 contigs < 200 kb. Outer ring (rainbow colors): Contigs and contig sizes. Outer track (Red/Black): MCIS coverage per sliding window. Window size = 1000 bp; Slide distance: 500 bp. Y-axes: average coverage per 1 kb window; axis limits set to min/max coverage. Inner track (Blue/Black): Repeat content per sliding window. Window size = 10 kbp; Slide distance: 5 kbp. Y-axes: repeat content in bp per 10 kb window; axis limits set to zero to maximum.

**S4 Fig.**
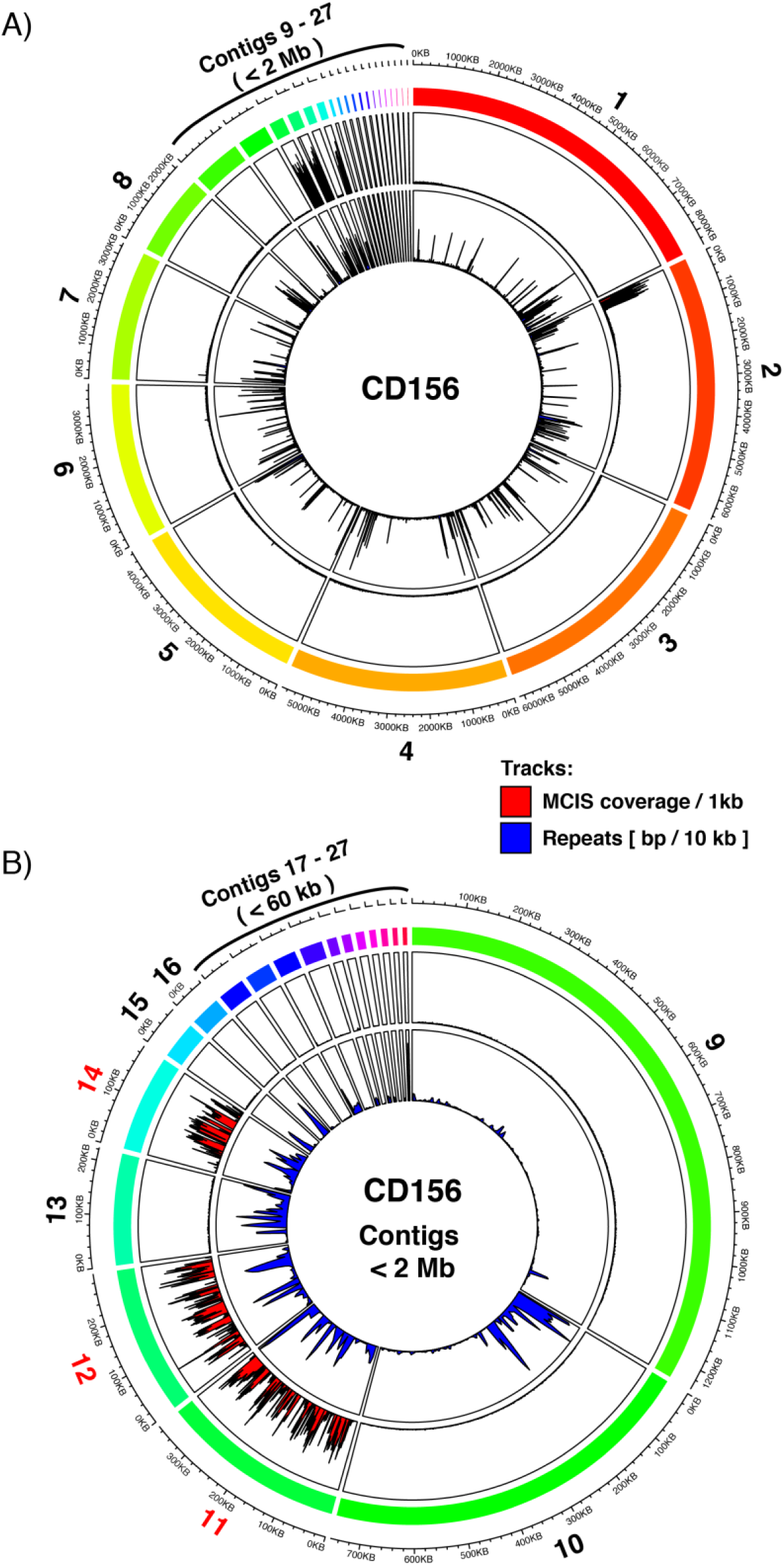
MCIS-read mapping and repeat content of *M. oryzae* strain CD156. **A)** Circos plot of mini-chromosome isolation sequencing (MCIS) coverage and repeat content across the CD156 genome. **B)** Circos plot of mini-chromosome isolation sequencing (MCIS) coverage and repeat content across US71 contigs < 2Mb. Outer ring (rainbow colors): Contigs and contig sizes. Outer track (Red/Black): MCIS coverage per sliding window. Window size = 1000 bp; Slide distance: 500 bp. Y-axes: average coverage per 1 kb window; axis limits set to min/max coverage. Inner track (Blue/Black): Repeat content per sliding window. Window size = 10 kbp; Slide distance: 5 kbp. Y-axes: repeat content in bp per 10 kb window; axis limits set to zero to maximum.

**S5 Fig.**
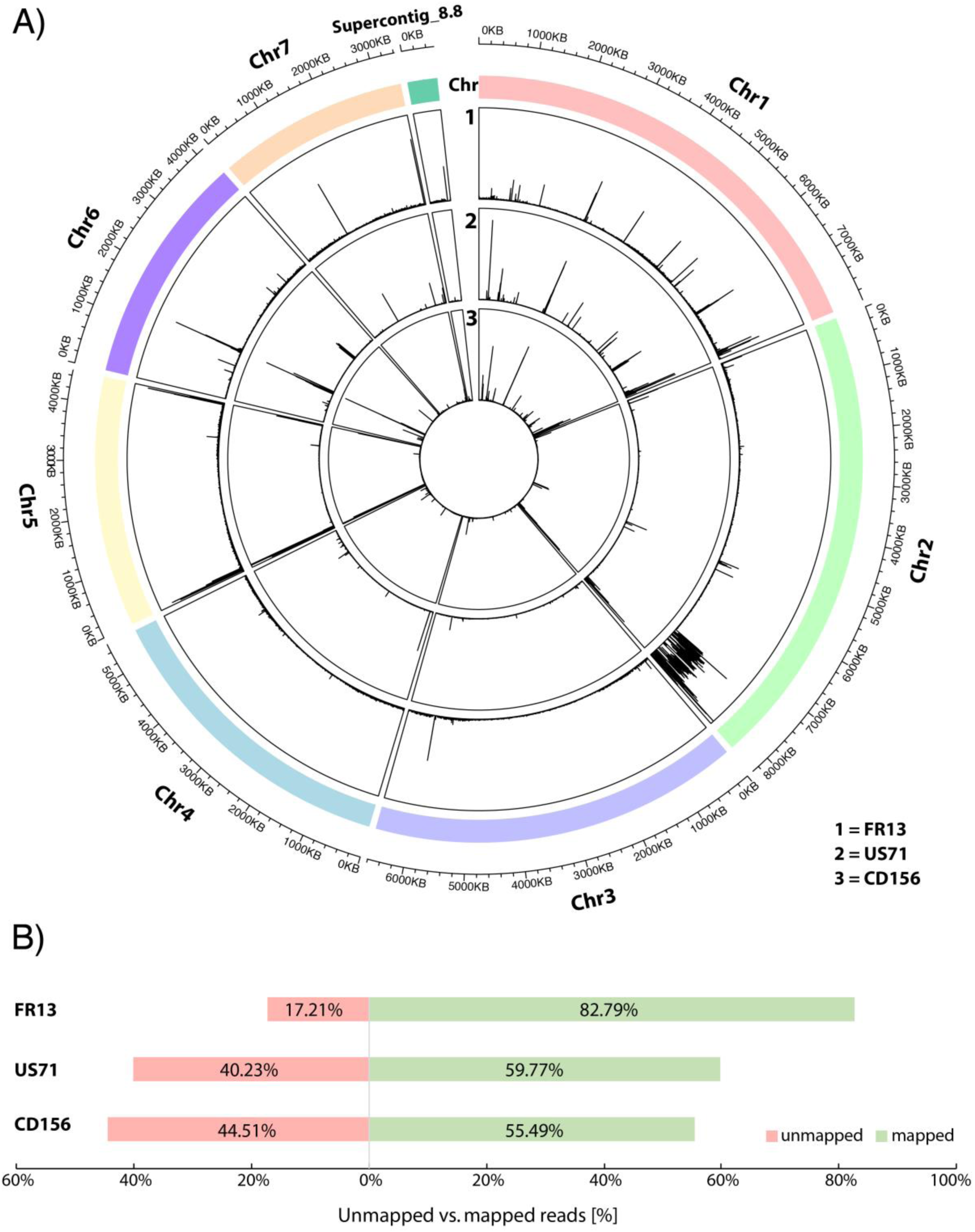
MCIS reads of FR13, US71, and US71 mapped against the reference genome of strain 70-15. **A)** Circos plot of MCIS reads uniquely mapped against the 70-15 genome. Outer ring: 70-15 chromosomes and chromosome sizes. Tracks: 1. FR13 MCIS read depth, 2. US71 MCIS depth, 3. CD156 MCIS read depth. **B)** Relative amount of MCIS total reads that mapped to the genome of strain 70-15. Mapped reads shown in green, unmapped reads in red.

**S6 Fig.**
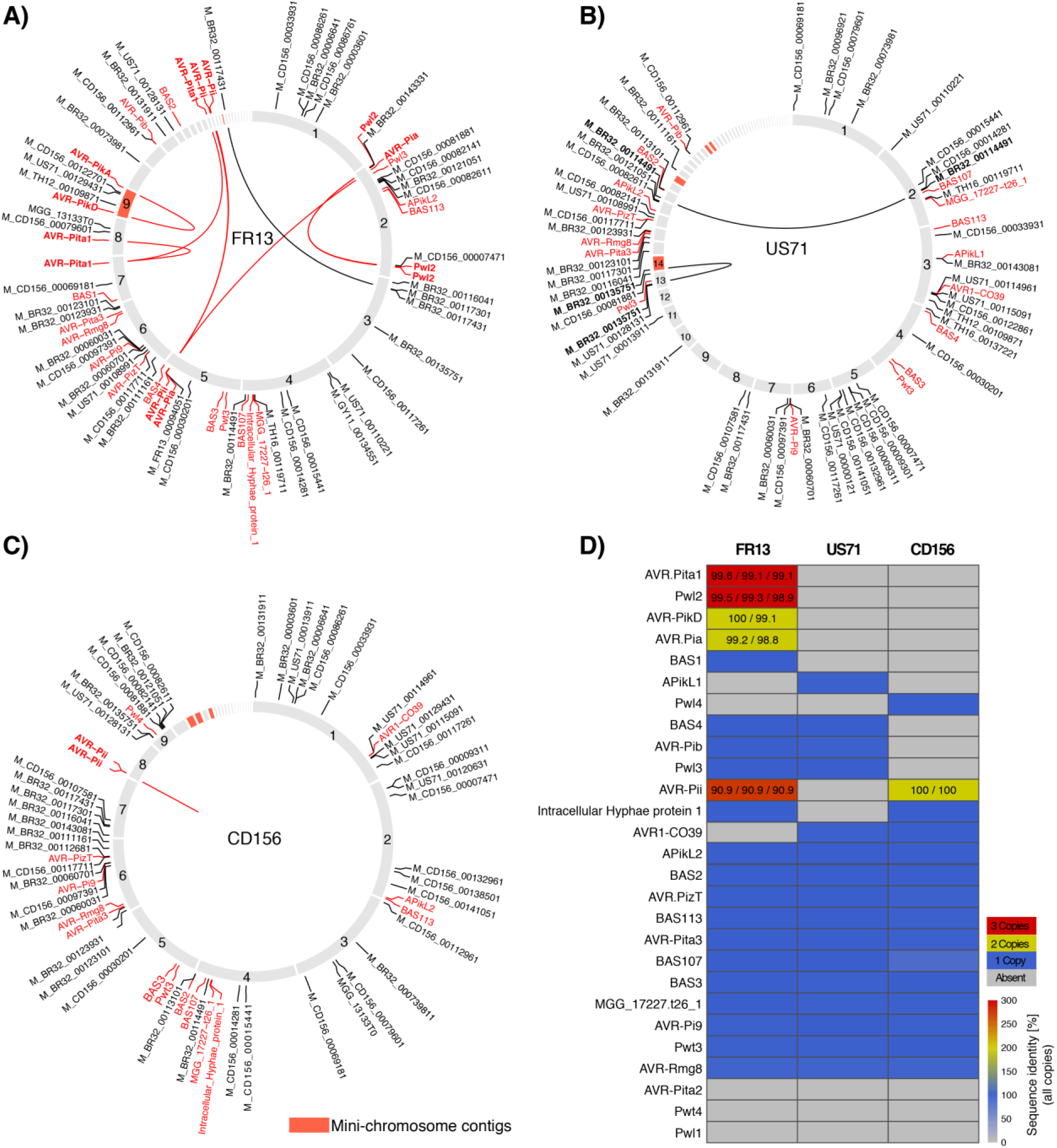
Known effector genes and predicted MAX-effectors are encoded on mini-chromosomes and can be duplicated between mini- and core-chromosomes. **A)** Position of effector genes in the FR13 genome. **B)** Position of effector genes in the US71 genome. **C)** Position of effector genes in the CD156 genome. Characterized effector genes are shown in red and predicted MAX-effectors are shown in black throughout A-C. Duplications are shown as lines in the center. Mini-chromosome contigs are shown in red, core-chromosomes in grey. **D)** Copy numbers of known effector genes in the genomes of isolates FR13, US71, and CD56. Absence shown in grey and presence shown in blue. Duplicated and triplicated genes are shown in yellow and red, respectively. Numbers in cells show the percentage identity of individual copies.

**S1 Table: Summary of gene and repeat content**

**S2 Table: Conservation and genomic location of orthologous groups**

**S3 Table: Summary of Pfam domain analysis**

**S4 Table: Summary of secreted protein prediction**

**S5 Table: Alignments and read support for merging of contigs S6 Table: Barcoding primers used for library preparation**

**S7 Table: Coverage of mini-chromosome derived reads and repeat content**

## References

1. Raffaele S, Win J, Cano LM, Kamoun S. Analyses of genome architecture and gene expression reveal novel candidate virulence factors in the secretome of *Phytophthora infestans*. BMC Genomics. 2010; 11:637.

2. Dong S, Raffaele S, Kamoun S. The two-speed genomes of filamentous pathogens: waltz with plants. Curr Opin Genet Dev. 2015; 35:57–65.

3. Croll D, McDonald BA. The Accessory Genome as a Cradle for Adaptive Evolution in Pathogens. PLoS Pathog. 2012; 8:e1002608.

4. Kämper J, Kahmann R, Bölker M, Ma L-J, Brefort T, Saville BJ, et al. Insights from the genome of the biotrophic fungal plant pathogen *Ustilago maydis*. Nature. 2006; 444:97–101.

5. Schirawski J, Mannhaupt G, Münch K, Brefort T, Schipper K, Doehlemann G, et al. Pathogenicity Determinants in Smut Fungi Revealed by Genome Comparison. Science. 2010; 330:1546–8.

6. De Jonge R, Bolton MD, Kombrink A, van den Berg GCM, Yadeta KA, Thomma BPHJ. Extensive chromosomal reshuffling drives evolution of virulence in an asexual pathogen. Genome Res. 2013; 23:1271–82.

7. Armitage AD, Taylor A, Sobczyk MK, Baxter L, Greenfield BPJ, Bates HJ, et al. Characterisation of pathogen-specific regions and novel effector candidates in *Fusarium oxysporum f. sp. cepae*. Sci Rep. 2018; 8:13530.

8. Bertazzoni S, Williams AH, Jones DA, Syme RA, Tan K-C, Hane JK. Accessories Make the Outfit: Accessory Chromosomes and Other Dispensable DNA Regions in Plant-Pathogenic Fungi. Mol Plant Microbe Interact. 2018; 31:779–88.

9. Goodwin SB, Ben M’Barek S, Dhillon B, Wittenberg AHJ, Crane CF, Hane JK, et al. Finished genome of the fungal wheat pathogen *Mycosphaerella graminicola* reveals dispensome structure, chromosome plasticity, and stealth pathogenesis. PLoS Genet. 2011; 7:e1002070.

10. Fokkens L, Shahi S, Connolly LR, Stam R, Schmidt SM, Smith KM, et al. The multi-speed genome of *Fusarium oxysporum* reveals association of histone modifications with sequence divergence and footprints of past horizontal chromosome transfer events. bioRxiv. doi: https://doi.org/10.1101/465070.

11. Chuma I, Isobe C, Hotta Y, Ibaragi K, Futamata N, Kusaba M, et al. Multiple Translocation of the AVR-Pita Effector Gene among Chromosomes of the Rice Blast Fungus *Magnaporthe oryzae* and Related Species. PLoS Pathog. 2011; 7:e1002147.

12. Frantzeskakis L, Kusch S, Panstruga R. The need for speed: compartmentalized genome evolution in filamentous phytopathogens: Multiple “speeds” in phytopathogen genomes. Mol Plant Pathol. 2019; 20:3–7.

13. Zolan ME. Chromosome-Length Polymorphism in Fungi. Microbiol Rev. 1995; 59:13.

14. Covert SF. Supernumerary chromosomes in filamentous fungi. Curr Genet. 1998; 33:311–9.

15. Jones RN, Viegas W, Houben A. A Century of B Chromosomes in Plants: So What? Ann Bot. 2008; 101:767–75.

16. Möller M, Stukenbrock EH. Evolution and genome architecture in fungal plant pathogens. Nat Rev Microbiol. 2017; 15:756–71.

17. Miao V, Covert S, VanEtten H. A fungal gene for antibiotic resistance on a dispensable (“B”) chromosome. Science. 1991; 254:1773–6.

18. Balesdent M-H, Fudal I, Ollivier B, Bally P, Grandaubert J, Eber F, et al. The dispensable chromosome of *Leptosphaeria maculans* shelters an effector gene conferring avirulence towards *Brassica rapa*. New Phytol. 2013; 198:887–98.

19. Ma L-J, van der Does HC, Borkovich KA, Coleman JJ, Daboussi M-J, Di Pietro A, et al. Comparative genomics reveals mobile pathogenicity chromosomes in *Fusarium*. Nature. 2010; 464:367–73.

20. Möller M, Habig M, Freitag M, Stukenbrock EH. Extraordinary Genome Instability and Widespread Chromosome Rearrangements During Vegetative Growth. Genetics. 2018; 210:517.

21. Fouché S, Plissonneau C, McDonald BA, Croll D. Meiosis Leads to Pervasive Copy-Number Variation and Distorted Inheritance of Accessory Chromosomes of the Wheat Pathogen *Zymoseptoria tritici*. Genome Biol Evol. 2018; 10:1416–29.

22. Habig M, Kema GH, Holtgrewe Stukenbrock E. Meiotic drive of female-inherited supernumerary chromosomes in a pathogenic fungus. eLife. 2018; 7:e40251. doi: 10.7554/eLife.40251

23. Raffaele S, Kamoun S. Genome evolution in filamentous plant pathogens: why bigger can be better. Nat Rev Microbiol. 2012; 10:417–30.

24. Coleman JJ, Rounsley SD, Rodriguez-Carres M, Kuo A, Wasmann CC, Grimwood J, et al. The Genome of *Nectria haematococca*: Contribution of Supernumerary Chromosomes to Gene Expansion. PLoS Genet. 2009; 5:e1000618.

25. Akagi Y, Taga M, Yamamoto M, Tsuge T, Fukumasa-Nakai Y, Otani H, et al. Chromosome constitution of hybrid strains constructed by protoplast fusion between the tomato and strawberry pathotypes of *Alternaria alternata*. J Gen Plant Pathol. 2009; 75:101–9.

26. Vanheule A, Audenaert K, Warris S, van de Geest H, Schijlen E, Höfte M, et al. Living apart together: crosstalk between the core and supernumerary genomes in a fungal plant pathogen. BMC Genomics. 2016; 17:670.

27. He C, Rusu AG, Poplawski AM, Irwin JA, Manners JM. Transfer of a supernumerary chromosome between vegetatively incompatible biotypes of the fungus *Colletotrichum gloeosporioides*. Genetics. 1998; 150:1459–66.

28. Fisher MC, Henk DanielA, Briggs CJ, Brownstein JS, Madoff LC, McCraw SL, et al. Emerging fungal threats to animal, plant and ecosystem health. Nature. 2012; 484:186–94.

29. Savary S, Willocquet L, Pethybridge SJ, Esker P, McRoberts N, Nelson A. The global burden of pathogens and pests on major food crops. Nat Ecol Evol. 2019; 3:430–9.

30. Hill L, Jones G, Atkinson N, Hector A, Hemery G, Brown N. The £15 billion cost of ash dieback in Britain. Curr Biol. 2019; 29:xR315–6.

31. Grünwald NJ, LeBoldus JM, Hamelin RC. Ecology and Evolution of the Sudden Oak Death Pathogen *Phytophthora ramorum*. Annu Rev Phytopathol. 2019; 57:301–21.

32. Białas A, Zess EK, De la Concepcion JC, Franceschetti M, Pennington HG, Yoshida K, et al. Lessons in Effector and NLR Biology of Plant-Microbe Systems. Mol Plant Microbe Interact. 2018; 31:34–45.

33. Lo Presti L, Lanver D, Schweizer G, Tanaka S, Liang L, Tollot M, et al. Fungal Effectors and Plant Susceptibility. Annu Rev Plant Biol. 2015; 66:513–45.

34. Kanzaki H, Yoshida K, Saitoh H, Fujisaki K, Hirabuchi A, Alaux L, et al. Arms race co-evolution of *Magnaporthe oryzae AVR-Pik* and rice *Pik* genes driven by their physical interactions. Plant J. 2012; 72:894–907.

35. Yoshida K, Saunders DGO, Mitsuoka C, Natsume S, Kosugi S, Saitoh H, et al. Host specialization of the blast fungus *Magnaporthe oryzae* is associated with dynamic gain and loss of genes linked to transposable elements. BMC Genomics. 2016; 17:370.

36. Jones JDG, Vance RE, Dangl JL. Intracellular innate immune surveillance devices in plants and animals. Science. 2016; 354:aaf6395–aaf6395.

37. Cesari S. Multiple strategies for pathogen perception by plant immune receptors. New Phytol. 2018; 219:17–24.

38. Dean R, Van Kan JAL, Pretorius ZA, Hammond-Kosack KE, Di Pietro A, Spanu PD, et al. The Top 10 fungal pathogens in molecular plant pathology: Top 10 fungal pathogens. Mol Plant Pathol. 2012; 13:414–30.

39. Ou SH. A look at worldwide rice blast disease control. Plant Disease. 1980; 64:439–45.

40. Kato H, Yamamoto M, Yamaguchi-Ozaki T, Kadouchi H, Iwamoto Y, Nakayashiki H, et al. Pathogenicity, mating ability and dna restriction fragment length polymorphisms of *Pyricularia* populations isolated from *Gramineae, Bambusideae* and *Zingiberaceae* Plants. J Gen Plant Pathol. 2000; 66:30–47.

41. Gladieux P, Condon B, Ravel S, Soanes D, Maciel JLN, Nhani A, et al. Gene Flow between Divergent Cereal- and Grass-Specific Lineages of the Rice Blast Fungus *Magnaporthe oryzae*. mBio. 2018; 9:e01219–17.

42. Gladieux P, Ravel S, Rieux A, Cros-Arteil S, Adreit H, Milazzo J, et al. Coexistence of Multiple Endemic and Pandemic Lineages of the Rice Blast Pathogen. Guttman D, editor. mBio. 2018; 9:e01806–17. https://doi.org/10.1128/mBio.01806-17.

43. Zhong Z, Chen M, Lin L, Han Y, Bao J, Tang W, et al. Population genomic analysis of the rice blast fungus reveals specific events associated with expansion of three main clades. ISME J. 2018; 12:1867–78.

44. Latorre SM, Reyes-Avila SC, Malmgren A, Win J, Kamoun S, Burbano HA. Recently expanded clonal lineages of the rice blast fungus display distinct patterns of presence/absence of effector genes. bioRxiv. 2020; Forthcoming.

45. Islam MT, Croll D, Gladieux P, Soanes DM, Persoons A, Bhattacharjee P, et al. Emergence of wheat blast in Bangladesh was caused by a South American lineage of *Magnaporthe oryzae*. BMC Biol. 2016; 14:84.

46. Islam MT, Kim K-H, Choi J. Wheat blast in Bangladesh: The current situation and future impacts. Plant Pathol J. 2019; 35:1–10.

47. Malaker PK, Barma NCD, Tiwari TP, Collis WJ, Duveiller E, Singh PK, et al. First report of wheat blast caused by *Magnaporthe oryzae* pathotype *triticum* in Bangladesh. Plant Dis. 2016; 100:2330–2330.

48. Chiapello H, Mallet L, Guérin C, Aguileta G, Amselem J, Kroj T, et al. Deciphering genome content and evolutionary relationships of isolates from the fungus *Magnaporthe oryzae* attacking different host plants. Genome Biol Evol. 2015; 7:2896–912.

49. Inoue Y, Vy TTP, Yoshida K, Asano H, Mitsuoka C, Asuke S, et al. Evolution of the wheat blast fungus through functional losses in a host specificity determinant. Science. 2017; 357:80–3.

50. Tosa Y, Osue J, Eto Y, Oh H-S, Nakayashiki H, Mayama S, et al. Evolution of an avirulence gene, AVR1-CO39, concomitant with the evolution and differentiation of *Magnaporthe oryzae*. Mol Plant Microbe Interact. 2005; 18:1148–60.

51. Stukenbrock EH. Evolution, selection and isolation: a genomic view of speciation in fungal plant pathogens. New Phytol. 2013; 199:895–907.

52. Talbot NJ, Salch YP, Ma M, Hamer JE. Karyotypic variation within clonal lineages of rice blast fungus *Magnaporthe grisea*. Appl Env Microbiol. 1993; 59:585–93.

53. Orbach MJ, Chumley FG, Valent B. Electrophoretic karyotypes of *Magnaporthe grisea* pathogens of diverse grasses. Mol Plant Microbe Interact. 1996; 9:261–71.

54. Luo C-X, Yin L-F, Ohtaka K, Kusaba M. The 1.6Mb chromosome carrying the avirulence gene AvrPik in *Magnaporthe oryzae* isolate 84R-62B is a chimera containing chromosome 1 sequences. Mycol Res. 2007; 111:232–9.

55. Kusaba M, Mochida T, Naridomi T, Fujita Y, Chuma I, Tosa Y. Loss of a 1.6 Mb chromosome in *Pyricularia oryzae* harboring two alleles of AvrPik leads to acquisition of virulence to rice cultivars containing resistance alleles at the Pik locus. Curr Genet. 2014; 60:315–25.

56. Peng Z, Oliveira-Garcia E, Lin G, Hu Y, Dalby M, Migeon P, et al. Effector gene reshuffling involves dispensable mini-chromosomes in the wheat blast fungus. Lin X, editor. PLOS Genet. 2019; 15:e1008272.

57. Win J, Chanclud E, Reyes-Avila S, Langner T, Islam MT, Kamoun S. Nanopore sequencing of genomic DNA from *Magnaporthe oryzae* isolates from different hosts. Zenodo. 2019. http://doi.org/10.5281/zenodo.2564950

58. Collemare J, Pianfetti M, Houlle A-E, Morin D, Camborde L, Gagey M-J, et al. *Magnaporthe grisea* avirulence gene *ACE1* belongs to an infection-specific gene cluster involved in secondary metabolism. New Phytol. 2008; 179:196–208.

59. De Guillen K, Ortiz-Vallejo D, Gracy J, Fournier E, Kroj T, Padilla A. Structure analysis uncovers a highly diverse but structurally conserved effector family in phytopathogenic fungi. PLOS Pathog. 2015; 11:e1005228.

60. Cruz CD, Valent B. Wheat blast disease: danger on the move. Trop Plant Pathol. 2017; 42:210–22.

61. Yadav V, Yang F, Reza MdH, Liu S, Valent B, Sanyal K, et al. Cellular dynamics and genomic identity of centromeres in cereal blast fungus. mBio. 2019; 10:e01581–19. doi: 10.1128/mBio.01581-192.

62. Faino L, Seidl MF, Shi-Kunne X, Pauper M, van den Berg GCM, Wittenberg AHJ, et al. Transposons passively and actively contribute to evolution of the two-speed genome of a fungal pathogen. Genome Res. 2016; 26:1091–100.

63. Depotter JRL, Shi-Kunne X, Missonnier H, Liu T, Faino L, van den Berg GCM, et al. Dynamic virulence-related regions of the plant pathogenic fungus *Verticillium dahliae* display enhanced sequence conservation. Mol Ecol. 2019; 28: 3482–3495. https://doi.org/10.1111/mec.15168

64. Fouché S, Badet T, Oggenfuss U, Plissonneau C, Francisco CS, Croll D. Stress-driven transposable element de-repression dynamics and virulence evolution in a fungal pathogen. Mol Biol Evol. 2019; msz216. https://doi.org/10.1093/molbev/msz216

65. Mehrabi R, Bahkali AH, Abd-Elsalam KA, Moslem M, Ben M’Barek S, Gohari AM, et al. Horizontal gene and chromosome transfer in plant pathogenic fungi affecting host range. FEMS Microbiol Rev. 2011; 35:542–54.

66. Langner T, Białas A, Kamoun S. The Blast Fungus Decoded: Genomes in Flux. mBio. 2018; 9:e00571–18. doi: 10.1128/mBio.00571-18

67. Schwessinger B. High quality DNA from Fungi for long read sequencing e.g. PacBio. protocols.io. 2016; doi: dx.doi.org/10.17504/protocols.io.ewtbfen.

68. Langner T, Harant A, Kamoun S. Isolation of supernumerary mini-chromosomes from fungi for enrichment sequencing. protocols.io. 2019; doi: dx.doi.org/10.17504/protocols.io.9t7h6rn

69. Koren S, Walenz BP, Berlin K, Miller JR, Bergman NH, Phillippy AM. Canu: scalable and accurate long-read assembly via adaptive *k* -mer weighting and repeat separation. Genome Res. 2017; 27:722–36.

70. Kurtz S, Phillippy A, Delcher AL, Smoot M, Shumway M, Antonescu C, et al. Versatile and open software for comparing large genomes. Genome Biol. 2004; 5, R12. doi: 10.1186/gb-2004-5-2-r12

71. Boetzer M, Henkel CV, Jansen HJ, Butler D, Pirovano W. Scaffolding pre-assembled contigs using SSPACE. Bioinformatics. 2011; 27:578–9.

72. Walker BJ, Abeel T, Shea T, Priest M, Abouelliel A, Sakthikumar S, et al. Pilon: An Integrated Tool for Comprehensive Microbial Variant Detection and Genome Assembly Improvement. PLoS ONE. 2014; 9:e112963.

73. Simão FA, Waterhouse RM, Ioannidis P, Kriventseva EV, Zdobnov EM. BUSCO: assessing genome assembly and annotation completeness with single-copy orthologs. Bioinformatics. 2015; 31:3210–2.

74. Bolger AM, Lohse M, Usadel B. Trimmomatic: a flexible trimmer for Illumina sequence data. Bioinformatics. 2014; 30:2114–20.

75. Li H, Handsaker B, Wysoker A, Fennell T, Ruan J, Homer N, et al. The Sequence Alignment/Map format and SAMtools. Bioinformatics. 2009; 25:2078–9.

76. Gu Z, Gu L, Eils R, Schlesner M, Brors B. circlize implements and enhances circular visualization in R. Bioinformatics. 2014; 30:2811–2.

77. Gel B, Serra E. karyoploteR: an R/Bioconductor package to plot customizable genomes displaying arbitrary data. Bioinformatics. 2017; 33:3088–90.

78. Altschul SF, Gish W, Miller W, Myers EW, Lipman DJ. Basic local alignment search tool. J Mol Biol. 1990; 215:403–10.

79. Nielsen H, Engelbrecht J, Brunak S, von Heijne G. Identification of prokaryotic and eukaryotic signal peptides and prediction of their cleavage sites. Protein Eng Des Sel. 1997; 10:1–6.

80. Emanuelsson O, Brunak S, von Heijne G, Nielsen H. Locating proteins in the cell using TargetP, SignalP and related tools. Nat Protoc. 2007; 2:953–71.

81. Krogh A, Larsson B, von Heijne G, Sonnhammer ELL. Predicting transmembrane protein topology with a hidden markov model: application to complete genomes. J Mol Biol. 2001; 305:567–80.

82. El-Gebali S, Mistry J, Bateman A, Eddy SR, Luciani A, Potter SC, et al. The Pfam protein families database in 2019. Nucleic Acids Res. 2019; 47:427–32.

83. Enright AJ. An efficient algorithm for large-scale detection of protein families. Nucleic Acids Res. 2002; 30:1575–84.

